# Multiomics reveal associations between CpG methylation, histone modifications and transcription in a species that has lost DNMT3, the Colorado potato beetle

**DOI:** 10.1101/2025.01.09.632173

**Authors:** Zoe M. Länger, Elisa Israel, Jan Engelhardt, Agata I. Kalita, Claudia I. Keller Valsecchi, Joachim Kurtz, Sonja J. Prohaska

**Author notes:** these authors contributed equally (Zoe M. Länger, Elisa Israel). these authors contributed equally (These authors jointly supervised this work) (Joachim Kurtz, Sonja J. Prohaska). E-Mail address of all authors: Zoe M. Länger, Elisa Israel, Jan Engelhardt, Agata I. Kalita, Claudia I. Keller Valsecchi,Sonja J. Prohaska.

## Abstract

Insects display exceptional phenotypic plasticity, which can be mediated by epigenetic modifications, including CpG methylation and histone modifications. In vertebrates, both are interlinked and CpG methylation is associated with gene repression. However, little is known about these regulatory systems in invertebrates, where CpG methylation is mainly restricted to gene bodies of transcriptionally active genes. A widely conserved mechanism involves the co-transcriptional deposition of H3K36 trimethylation and the targeted methylation of unmethylated CpGs by the *de novo* DNA methyltransferase DNMT3. However, DNMT3 has been lost multiple times in invertebrate lineages raising the question of how the links between CpG methylation, histone modifications and gene expression are affected by its loss.

Here, we report the epigenetic landscape of *Leptinotarsa decemlineata,* a beetle species that has lost DNMT3 but retained CpG methylation. We combine RNA-seq, enzymatic methyl-seq and CUT&Tag to study CpG methylation and patterns of H3K36me3 and H3K27ac histone modifications on a genome-wide scale. Despite the loss of DNMT3, H3K36me3 mirrors CpG methylation patterns. Together, they give rise to signature profiles for expressed and non-expressed genes. H3K27ac patterns, which show no association with CpG methylation, have a prominent peak at the transcription start site that is predictive of expressed genes. Our study provides new insights into the evolutionary flexibility of epigenetic modification systems that urge caution when generalizing across species.

**Research highlights:** Despite lacking DNMT3, EM-seq revealed CpG methylation in the Colorado potato beetle. CUT&Tag showed an association of H3K36me3 and H3K27ac with transcription, while only H3K36me3 aligns with CpG methylation, demonstrating epigenetic flexibility.

## Introduction

The remarkable evolutionary success of insects is largely due to their exceptional ability to generate phenotypic diversity. The basis for such plasticity is partially provided by epigenetic modifications, which facilitate differential gene expression. This is achieved through reversible chemical modifications on histones or DNA bases (Aliaga et al., 2019). While epigenetic modifications as such are well conserved, their functions and interconnections might be evolutionarily flexible. However, we currently know little about this beyond a few intensively studied model organisms.

The most prominent DNA methylation involves the addition of a methyl group to position 5 of cytosine in a CpG dinucleotide context (Lister et al., 2009). This modification is established *de novo* by DNA methyltransferase 3 (DNMT3) and faithfully maintained by DNMT1 during DNA replication (Lyko, 2017). In vertebrates, CpG methylation is present at medium to high levels and is mainly associated with gene silencing (Bourc’his & Bestor, 2004; Greenberg & Bourc’his, 2019). Conversely, in invertebrates, CpG methylation predominantly occurs in gene bodies, especially in exons of active genes as reported for multiple species, including the honey bee (*Apis mellifera*) and the silk moth (*Bombyx mori*) (Lewis et al., 2020; Suzuki et al., 2007; Zemach et al., 2010). However, changes in methylation do not seem to lead to coordinated changes in transcription (Cardoso-Júnior et al., 2021; Dixon & Matz, 2022; Harris et al., 2019).

Many invertebrate model organisms, such as *Drosophila melanogaster* and *Caenorhabditis elegans*, lack detectable levels of CpG methylation (Raddatz et al., 2013; Zemach et al., 2010), reflecting the recurrent loss of DNA methylation throughout Ecdysozoan evolution (Engelhardt et al., 2022), and casting doubt on the central role of CpG methylation in gene regulation. In Coleoptera, some species have lost CpG methylation (e.g. *Tribolium castaneum* (Schulz et al., 2018)), while others, like *Nicrophorus vespilloides*, retained low levels (Cunningham et al., 2015). The focal association of gene body methylation with actively transcribed genes and the overall reduction of CpG methylation in holometabolous insects suggest divergent functional evolution of CpG methylation in vertebrates and invertebrates (Hunt et al., 2013; Provataris et al., 2018). Many species with low CpG methylation have lost DNMT3 but retained DNMT1 (Engelhardt et al., 2022). This may suggest that DNMT1 alone is maintaining CpG methylation over many generations, or acts as a *de novo* methyltransferase under certain conditions (Glastad et al., 2019; Dahlet et al., 2020).

Modifications on histones can alter the structure of chromatin and affect various parts of the gene transcription process, which can lead to correlations between histone modification and gene expression (Zhang et al., 2022). A core set of histone modifications is broadly conserved across eukaryotes, including H3 and H4 lysine acetylation marks, indicative of gene expression-permissive chromatin and H3 lysine trimethylation marks, which demarcate either active (e.g., H3K4me3 and H3K36me3) or repressive (e.g., H3K27me3 and H3K9me3) chromatin states (Grau-Bové et al., 2022). In particular, H3K36me3 is prevalent on gene bodies of active genes and associated with transcription (Guenther et al., 2007; Pokholok et al., 2005). In contrast, acetylation of histone H3 lysine 27 (H3K27ac) is commonly associated with regulatory regions, i.e. promoters and enhancers, in the context of gene activity (Nègre et al., 2011; Simola et al., 2013). H3K27ac is involved in nucleosome mobilization and eviction (Kang et al., 2021), at the transcription start site (TSS) of active genes. Furthermore, H3K27ac, set by homologs of the histone acetyltransferase p300/CBP, antagonizes the establishment of H3K27me3, the repressive mark set by polycomb complex 2 (PRC2).

In vertebrates, DNA methylation and histone modification are highly interlinked. An association of CpG methylation and H3K36 methylation appears to be deeply conserved among eukaryotes (de Mendoza et al., 2020; Hahn et al., 2011; Neri et al., 2017; Pokholok et al., 2005; Yano et al., 2022), but studies in insects showing this association are limited. Gene body methylation in *A. mellifera* and *Solenopsis invicta* (fire ant) positively correlates with activating histone marks in *D. melanogaster,* e.g. H3K36me3 (Hunt et al., 2013; Nanty et al., 2011). A link between H3K27ac and CpG methylation, mediated by the methyl-CpG-binding domain protein 2/3 (MBD2/3), was described by (Xu et al., 2021) in *B. mori*. The authors propose that MBD2/3 targets Tip60, a histone H3K27 acetyltransferase complex, to methylated CpGs.

DNMTs contain domains for recognition of histone modification marks and, *vice versa*, histone modifying complexes and transcription factors distinguish between methylated and unmethylated CpGs. While maintenance of CpG methylation by DNMT1 is coupled to DNA replication, *de novo* establishment by DNMT3 is promoted by transcription. Here, SETD2 interacts with the elongating RNA-Pol II to deposit H3K36me3 at the gene body (Edmunds et al., 2007; Yoh et al., 2008). In turn, DNMT3 binds to H3K36me3 via its PWWP domain and methylates unmethylated CpGs (Dhayalan et al., 2010; Teissandier & Bourc’his, 2017; Wagner & Carpenter, 2012). The functional overlap between CpG methylation and other epigenetic mechanisms, such as certain histone modifications, may range from perfect complementarity – rendering methylation indispensable – to full redundancy, which would facilitate the loss of CpG methylation (Hunt et al., 2013).

For species that lost DNMT3, the question arises of how CpG methylation marks can be set to genes that become active later in development. Experimental data on DNA methylation in Coleoptera is currently only available for two species, *T. castaneum* (loss of DNMT3 and DNA methylation) and *N. vespilloides* (DNMT1, DNMT3 and DNA methylation present) (Cunningham et al., 2015; Schulz et al., 2018). We thus studied the Colorado potato beetle (*L. decemlineata)* as the first beetle species, where DNA methylation was predicted to be retained despite the loss of DNMT3 (Engelhardt et al., 2022).

This study aims to expand our knowledge of the organization of epigenetic systems in insects, as an in-depth understanding of the underlying mechanisms is needed to appraise the epigenetic contribution to phenotypic diversity and evolutionary novelty (Maleszka, 2024). Here, we experimentally assess and compare CpG methylation of *L. decemlineata* in adults and embryos, the genomic location of H3K36me3 and H3K27ac in embryos, and link that epigenetic information to genome-wide gene expression data. We combined RNA sequencing with novel techniques for studying CpG methylation and histone modifications, enzymatic methyl-seq (EM-seq) and CUT&Tag, respectively, demonstrating the potential of whole-genome approaches even in species whose epigenomes were previously largely unstudied.

## Material and Methods

### Model organism and samples

We used Colorado potato beetles, *L. decemlineata* (Coleoptera). For detailed rearing conditions see supplementary methods. For replicates of embryo EM-seq, RNA-seq and CUT&Tag, we pooled approximately 30 embryos from different egg clusters laid on the same day. For adult sample preparation, we pooled two individuals per replicate. We sampled the living animals of approximately the same age (2-3 weeks after eclosion) on the same day for sequencing and stored them at -80°C until processing, if necessary. For EM-seq and RNA-seq, we used 3 replicates for each life stage (mixed sex). For CUT&Tag, we used 2 embryo replicates.

We used the chromosome-level genome assembly and gene annotation of *L. decemlineata* provided by the Gene Expression Atlas of the Colorado Potato Beetle, denoted as version LdNA_01 (Wilhelm et al., 2024). The assembly is identical to ASM2471293v1/GCA_024712935.1 (Yan et al., 2023) on NCBI/GenBank, respectively.

### DNA methylation: Enzymatic methyl sequencing (EM-seq)

High molecular weight DNA was extracted from pooled whole body embryo or adult *L. decemlineata* using a combination of chloroform:isoamyl and a salting out procedure. Detailed protocol information can be found in the supplementary methods. Enzymatic methyl library preparation and sequencing were kindly performed in the Cologne Center for Genomics (CCG) University of Cologne, Cologne, Germany. The enzymatic conversion method (EM-seq) was used, as described in (Vaisvila et al., 2021). Methylated (CpG methylated pUC19) and unmethylated (unmethylated Lambda) controls were included in the library preparation and we checked the conversion rate for these controls after sequencing. Briefly, 200 ng high molecular weight DNA was processed using NEBNext UltraShear, following the manufacturer’s instructions. For a fragment length of 250-350 bp, Enzymatic fragmentation was conducted for 20 minutes at 37°C and 15 minutes at 65°C to hold at 4°C. For PCR library amplification, the following program was used: (1. 30“-98°C, 2. 10“-98 °C, 3. 30“-62 °C, 4. 1’-65 °C (2.-4. ×4), 5. 5-65 °C). Amplified libraries were cleaned with 65µL of resuspended NEBNext Sample Purification Beads. Libraries were sequenced on an Illumina NextSeq 2000 (2 x 150 bases).

#### Data processing (EM-seq)

We trimmed the reads using trim_galore (Krueger et al., 2023; Martin, 2011) in paired-end mode. We used default settings but trimmed 10 nucleotides from both sides of each read (–clip_R1=10, –clip_R2=10, –three_prime_clip_R1=10, –three_prime_clip_R2=10) to remove potential remnants of EM-seq adapters. Subsequently, the reads were mapped, deduplicated and the methylation level extracted using the Bismark pipeline (Krueger & Andrews, 2011). The number of reads and coverage can be found in Tab. S1. To accommodate the lower quality of the Coleoptera assembly compared to vertebrate model organisms we used a lower minimum alignment score (--score_min L,0,-0.6). The enrichment of methylation levels in different parts of the genome was calculated using BEDTools (Krueger & Andrews, 2011; Quinlan & Hall, 2010) and custom shell scripts. To estimate CpG methylation levels, we require cytosines (in a CpG context) to be covered by five or more reads. We then define the methylation level of a cytosine as the fraction of the number of methylated reads divided by the total number of reads covering that cytosine. To estimate the degree of methylation of a genomic feature, such as a gene, we calculate the mean percent methylation of all CpGs within that feature, if and only if the feature contains three or more CpGs. The consolidated set of genes is generated by excluding all genes that have less than three CpGs per gene in one or more of the six samples (3x embryo and 3x adult). All replicates correlate strongly (Pearson, p < 0.05) regarding all individual CpG sites as well as in the methylation levels of genes (see Fig. S1-4). We used RepeatModeller v2.0.5 (Flynn et al., 2020) to generate models of de novo repeats and RepeatMasker v4.1.7-p1 to annotate them in the genome.

We used metilene version 0.2-8 (Jühling et al., 2016) to predict differentially methylated regions (DMRs) between the adult and embryo replicates. Only CpGs with 5 or more and less than or equal to 100 reads in all six replicates were used as input. A DMR was considered significant if it had a Benjamini-Hochberg adjusted p-value <0.05. The resulting DMRs have at least 10 CpGs and a mean methylation difference of at least 10%.

### Gene expression: RNA extraction and RNA-seq

Pooled whole-body samples of embryos or adults were taken for RNA extraction. We used a protocol combining Trizol lysis and chloroform extraction with the purification via spin columns from the SV Total RNA Isolation System (Promega). The detailed protocol can be found in the supplementary methods.

#### Data processing (RNA-seq)

RNA sequencing and basic data processing was carried out by Novogene (Planneg, Munich, Germany). In short samples were sequenced using the Illumina NovaSeq PE150 platform. The raw data was cleaned to remove reads containing adapters, more than 10% N’s or when low quality nucleotides constituted more than 50% of the read. The number of reads and coverage can be found in Tab. S2. Subsequently, HISAT2 version 2.0.5 (Mortazavi et al., 2008) was used to align the reads to the genome and DESeq2 version 1.20.0 (Love et al., 2014) with Benjamin Hochberg adjusted p-value to predict differentially expressed genes.

One embryonic replicate (embryo 1) did neither correlate (Pearson correlation) nor cluster (Principal component analysis) with the other embryonic or adult replicates (see Fig. S5-7). Thus, we dismissed it in further analyses and rerun DESeq2, using NovoMagic, a free Novogene platform for data analysis.

### Histone modifications: CUT&Tag library generation and sequencing

We performed CUT&Tag as described in (Kaya-Okur et al., 2020) with slight modifications. Whole-body samples of the studied species were flash-frozen, homogenized in cold DPBS (supplemented with MgCl2 and CaCl2), and passed through a cell strainer (Corning, 352235). We used 0.4 million cells per reaction from whole-body samples of the studied species. Cells were bound to ConA beads in a 1:10 ratio for 10 min at room temperature. We then incubated the cells in an antibody buffer with the primary antibody (1:100) [IgG control Rabbit (Abcam catalog no. ab37415), H3K36me3 (Active Motif catalog no. 91266), H3K27ac (Abcam catalog no. ab4729)] at 4°C overnight on a nutator, which was followed by incubation with the secondary antibodies (1:100) [αMs IgG Rabbit (Abcam catalog no. ab6709), αRb IgG Guinea pig (Sigma-Aldrich catalog no. SAB3700890)] for 60 min at room temperature. Cells were rinsed, washed twice, and incubated with loaded pA-Tn5 (1:200) for 1 h at room temperature on a nutator. To remove excess pA-Tn5, cells were rinsed and then washed. To perform the tagmentation, we incubated the cells with 10mM MgCl_2_ for 1 h at 37°C. To stop the tagmentation and solubilize the DNA fragments, we added EDTA, SDS and Proteinase K and incubated for 1h at 55°C. Libraries were amplified with the NEBNext Ultra Q5 Master Mix (NEB) in 14 cycles and purified using the DNA Clean & Concentrator-5 Zymo kit, following the manufacturer’s instructions. Amplified libraries were resuspended in 15µL nuclease free H_2_O. We quantified library concentrations using Qubit and measured the library size on Bioanalyzer.

Pooled samples were sequenced on Illumina NextSeq 500 High Output, PE for 2×75 cycles plus 2×8 cycles for the dual index read by the Institute for Molecular Biology, Mainz, Genomics Core Facility. pA–Tn5 was prepared by the Institute for Molecular Biology, Mainz, Protein Production Core Facility.

#### Data processing (CUT&Tag)

We removed adapters using cutadapt version 4.8 (Martin, 2011). With bowtie2 (version 2.5.3) (Langmead & Salzberg, 2012), we aligned the trimmed reads to the reference genome of *L. decemlineata*, with the following parameters --end-to-end --very-sensitive --no-mixed --no-discordant --phred33 -I 10 -X 700. Given that we did not expect PCR duplication, no duplicates were removed. The number of reads and alignment rate can be found in the Tab. S3, fragment length in Fig. S8. We calculated a Pearson correlation between the replicates and samples (see Fig. S9) with a custom R script. To assess the coverage of features, we employed bedtools (version 2.31.1) genomecov (Quinlan & Hall, 2010). For peak calling and sparse enrichment analysis, we used SEACR version 1.3 (Meers et al., 2019) with parameters set to ‘norm’ and ‘stringent’. SEACR was given the parameter ‘0.025’ in order to obtain the 2.5% of highest peaks for each replicate. Subsequently, we examined the sets for overlaps using bedtools intersect and merged the resulting subset of overlapping peaks with bedtools merge. This way, we obtained a high-confidence set of reproducible peaks. We visualized heatmaps using deepTools version 3.5.5 (Ramírez et al., 2016), employing the functions computeMatrix --scale-regions and plotHeatmap. For this, we normalized the gene length to a length of 5 kb, with 3 kb upstream and downstream of the gene body.

### Gene Ontology (GO) enrichment analysis

Gene Ontology (GO) enrichment analysis was conducted using the topGO package version 2.56.0 (Adrian Alexa, 2017) in R. We used the Parent-Child algorithm to determine the highest common GO term level and identified significantly enriched GO terms using the classic Fisher’s exact and corrected the p-value for multiple tests using the Benjamini-Hochberg method. Gene ratio was calculated as the percentage of genes annotated with the respective GO term in a subset to the entity of genes with the respective GO term in the genome. The 40 most significant GO terms were visualized using ggplot2 (Wickham et al., 2007).

### Data visualization

Part of the data was analyzed and plotted using R version 4.4.1 (R Core team, 2024) and RStudio (R Studio team, 2024). Data manipulation (includes filtering, summation and reshaping) was performed using dplyr version 1.1.4 (Wickham, François, et al., 2014), tidyr version 1.3.1 (Wickham, Vaughan, et al., 2014) and Hmisc version 5.1-3 (Harrell, 2003), also using reshape2 (Wickham, 2007) and ggExtra version 0.10.1 (Attali & Baker, 2015). Scatterplots and density plots were generated using ggplot2 (Wickham et al., 2007), Venn diagrams with VennDiagram version 1.7.3 (Chen, 2011) and the Sankey plot using ggalluvial version 0.12.5 (Brunson & Read, 2017).

## Results

### CpG methylation is highest in exons

To quantify CpG methylation, we performed EM-seq on three replicates each of embryo and adult samples of *L. decemlineata*. On a genome-wide level, 4.2% and 3.4% of all CpGs were methylated in embryos and adults, respectively. Among genomic features, exons show the highest mean percent methylation with 16.5% and 13.8% in embryos and adults, respectively, followed by introns, while intergenic regions have very low levels of methylation (Fig. 1A, Tab. S4-5). The seemingly large number of CpGs in introns can be explained by the different total size of intron and exon sequences in the genome (Fig. 1A). Of the entire *L. decemlineata* genome, appr. 67% was repeat masked, which is similar to (Yan et al., 2023). Transposable elements (LINEs, LTR elements, DNA transposons) cover appr. 41% of the genome and have a methylation level of 3.2%. The remaining repeats are almost exclusively unclassified.

**Figure 1.**
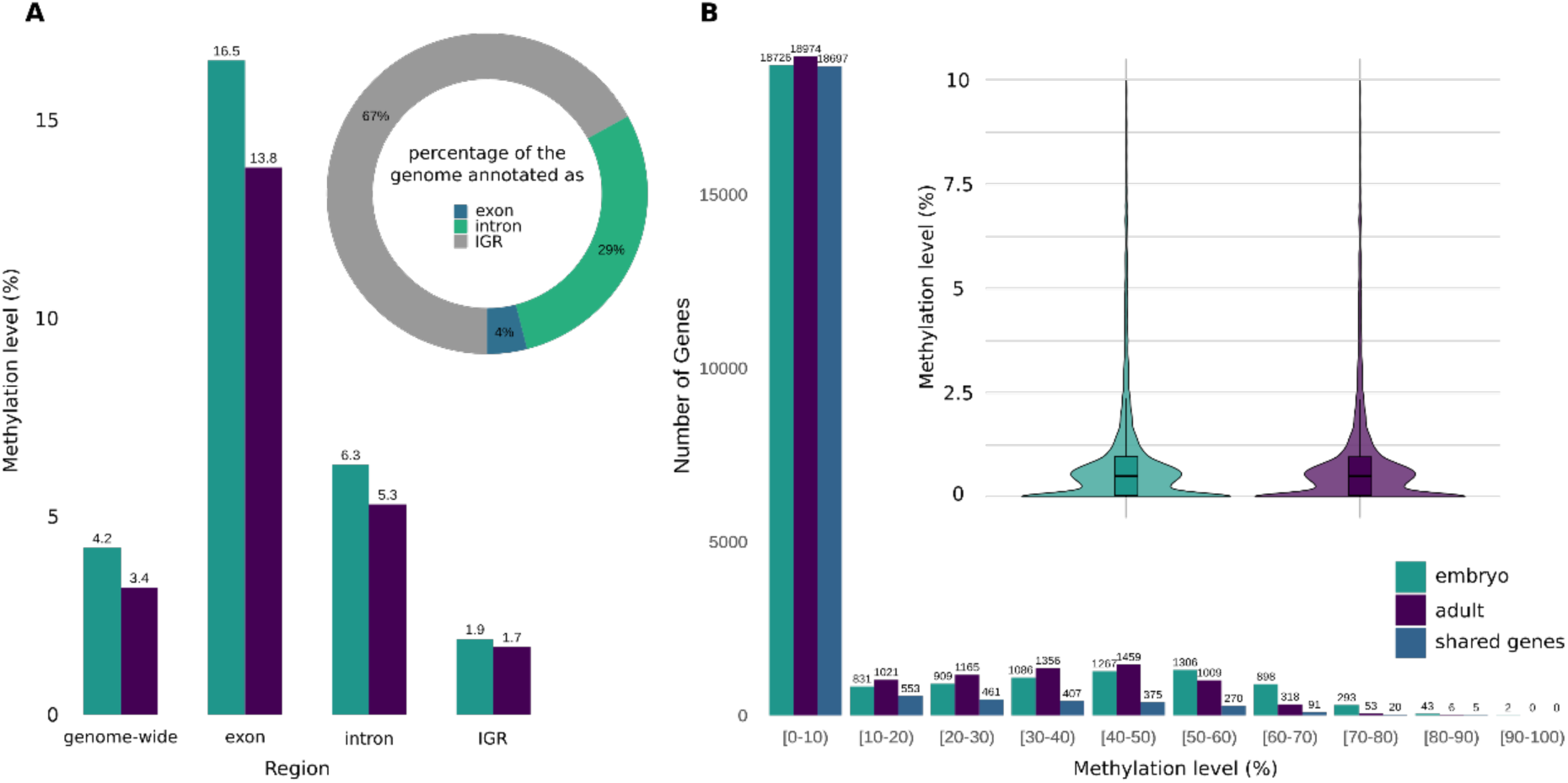
Distribution of mean CpG methylation levels. **A) across different genomic regions and B) across genes, in embryos and adults.** Inset in A: percentage of the genome annotated as exon, intron or intergenic region. Inset in B: zoom onto the distribution of genes in the [0-10) interval.

Focusing on genes, the majority of genes are not methylated (below 10%). Most of the methylated genes show mean methylation levels up to 70% while only 1.5% of genes in embryos and 0.3% in adults show methylation levels above 70% (Fig. 1B).

### Dividing annotated genes into four subsets based on methylation and expression status

To study the relationship between gene expression, gene body methylation and histone modifications, we generated RNA-seq data for embryo and adult samples. Comparing embryo and adult gene expression, we identified 5,135 differentially expressed genes; 3,790 with higher expression in adults and 1,345 genes with higher expression in embryos (Fig. S10). From now on, a differentially expressed gene (DEG) will be referred to as “downregulated” if the expression is higher in the embryo compared to the adult, otherwise as “upregulated”.

Due to the pronounced level of methylation in genes, we narrowed our analysis down to a consolidated set of 25,631 protein-coding genes. In this set, all genes had sufficient coverage of DNA methylation and RNA-seq data for all replicates. We then assigned genes to the categories *’not methylated’*, i.e. methylation level below 10% vs. *’methylated’*; and *’not expressed’*, i.e. FPKM 1 or below *vs. ‘expressed’*. In combination, this results in four groups of genes, namely, genes which are *’not methylated / expressed’*, *‘methylated / expressed’*, *’not methylated / not expressed’*, and *’methylated / not expressed’*. In the following, all our analyses will refer to these four mutually exclusive gene sets if not stated otherwise. The biggest set is *’not methylated / not expressed’* and the smallest set is *’methylated / not expressed’.* The number of genes in each set for embryo and adult can be found in Tab. S7.

### *‘Methylated / expressed*’ genes are usually longer and exclusively characterized by a drop in CpG methylation at the TSS

We found *‘methylated / expressed’* genes in both embryo and adult exhibit the longest mean length (appr. 2 kb), while *‘not methylated / not expressed’* genes displayed the shortest mean lengths (720 bp and 820 bp, respectively), see Fig. 2 (Fig. S11 for adults). Certain subsets displayed considerable variability in their gene length, highest in *‘methylated / not expressed’* genes and lowest in *‘not methylated / not expressed’* genes, see Tab. S8.

**Figure 2.**
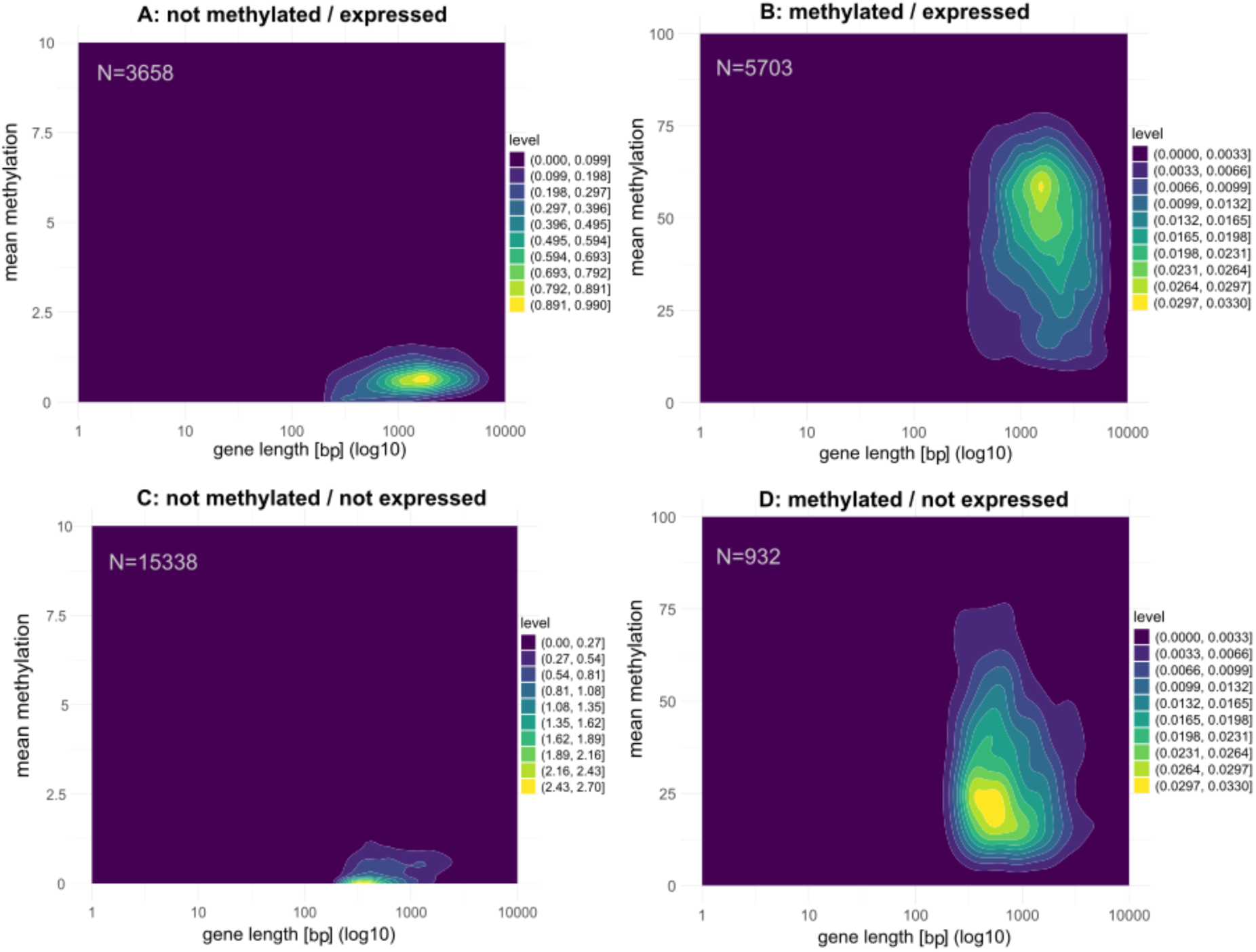
Relationship between gene length and mean methylation (%) in different categories. (Note difference in scale). The color gradient (level) represents the density of genes at different methylation levels, with brighter shades indicating regions of higher density.

We examined the CpG methylation of the gene body as well as the 3kb upstream and downstream regions in embryos and adults. In general, exon methylation exceeds intron methylation as described in previous studies (Lewis et al., 2020), and gene body methylation is more pronounced than methylation of non-genic flanking regions (see Fig. S12, Tab. S9, S10). Surprisingly, we observe a prominent drop in CpG methylation at the TSS in *‘methylated / expressed’* genes that is clearly absent in *‘methylated / not expressed’* genes. Furthermore, CpG methylation levels in the downstream regions decrease only slowly with greater distance from the transcription end site (Fig. 3).

**Figure 3.**
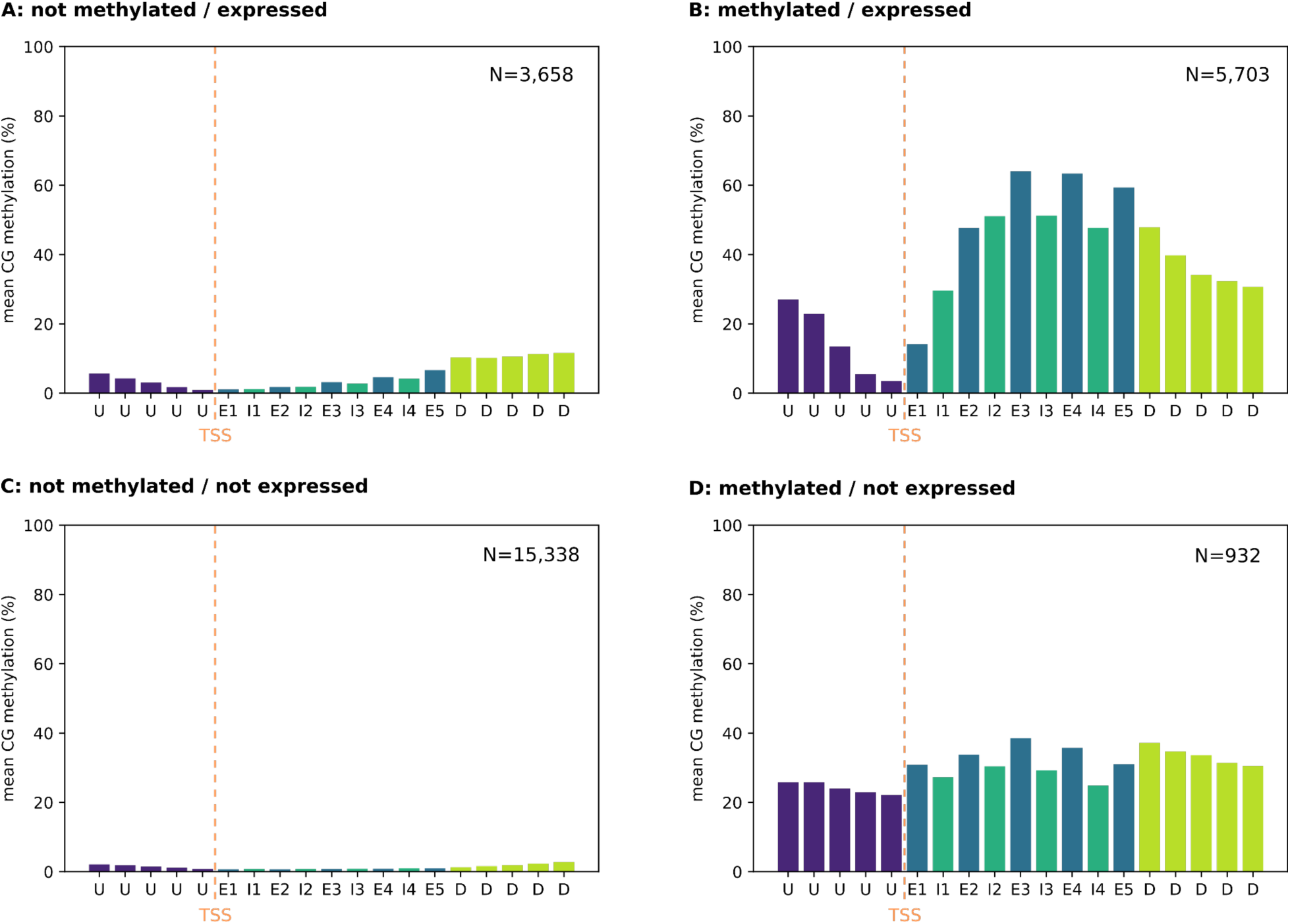
Mean embryonic methylation level of different genic segments. E1 to E5 represent the first 5 exons; I1-I4 represent the first four introns; U – upstream region; D – downstream region.

### Gene body methylation is associated with transcription, but changes in GBM are not associated with transcription changes

We identified 3,822 significantly differentially methylated regions (DMRs) between the adult and embryo stage. 3,754 DMRs have a lower methylation in adults compared to embryos, we denote them hypomethylated. 68 DMRs are hypermethylated in the adult stage. Most hypo- and about half of the hypermethylated DMRs overlap with genes (including 3kb up- and downstream) while only 11% of the hypo- but 51% of the hypermethylated DMRs are located in the intergenic region.

Of the hypomethylated DMRs that overlap genes, 1,469 overlap only one genic feature; similarly for 31 hypermethylated DMRs. For hypomethylated DMRs overlapping more than one genic feature the most frequent class are 1,152 DMRs which overlap both exons and intron (see Fig. 4).

**Figure 4.**
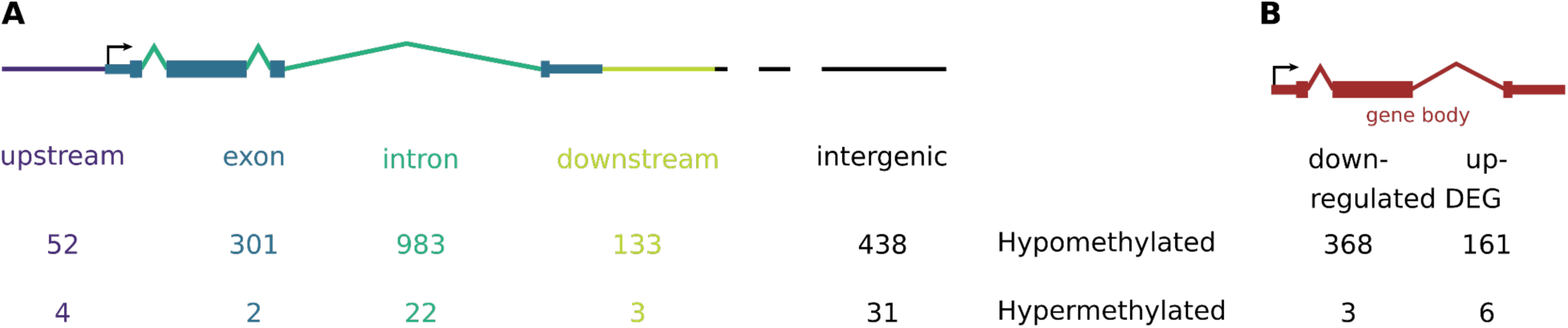
**A) Differentially methylated regions (DMR) per genic feature.** Shown are the numbers of differentially methylated regions overlapping exactly one genic feature, using the embryo stage as reference. Some regions overlap with multiple genic regions. **B) Relationship between differentially methylated genes and differentially expressed genes (DEGs).**

Exploring the association between gene body methylation and gene expression we found a weak but significant positive linear correlation in ‘*methylated / expressed’* genes for both embryos and adults (Fig. 5A; Fig. S13 for adults). However, methylation explains only a small portion of the variability in expression, and neither linear nor quadratic regression models account for much of the total variation (see Fig. S14). We cannot determine whether methylation promotes gene expression or v*ice versa*. A matrix summarizing ‘*methylated’* and ‘*not methylated’ genes,* categorized by their expression status (‘*expressed’/’not expressed’*), is provided in Fig. S15.

**Figure 5.**
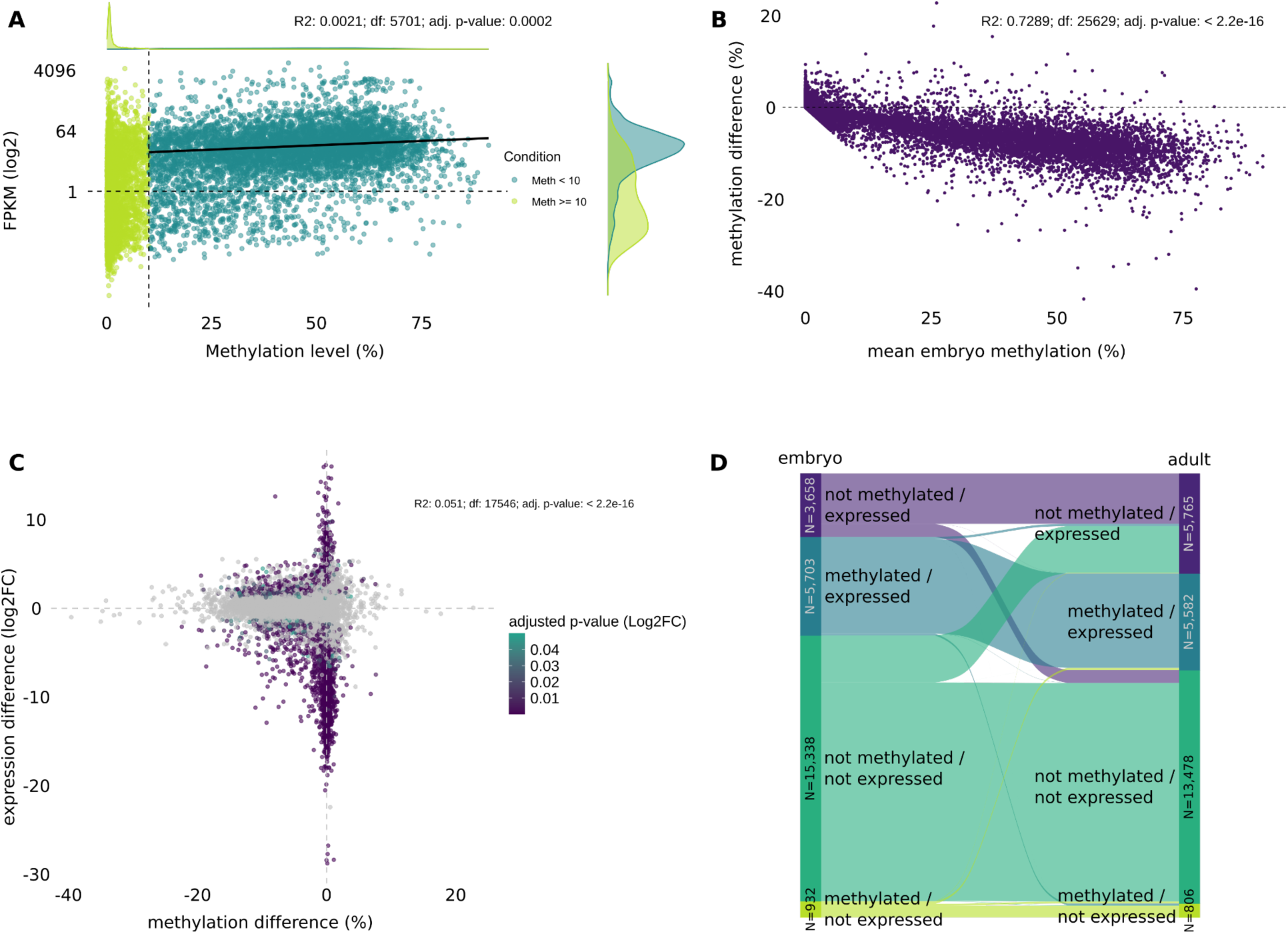
Changes in gene body methylation and gene expression. **A) Association between gene expression level (FPKM) and gene body methylation level in embryos.** Dashed gray lines indicate significance thresholds. All values that equal 0 were removed and linear regression was calculated on the subset of methylated/expressed genes **B) Observed changes in gene body methylation from embryo to adult relative to embryonic methylation levels. C) Relationship of gene body methylation and gene expression between embryo and adult.** Shown is the correlation of differences in gene body methylation and gene expression (log_2_ fold change [Log_2_FC]) between embryonic and adult life stages. Colored dots indicate significant changes in gene expression. Grey points indicate non-significant changes in Log_2_FC. Pearson Correlation coefficient was calculated between the methylation difference and the expression difference. **D)Sankey diagram showing the change of gene expression and methylation between embryo and adult stage.**

We examined changes in gene expression and CpG methylation accompanying the transition from embryo to adult between the four categories (Fig. 5D). 9,361 (36.5%) and 11,347 (44.3%) genes are expressed in embryos and adults, respectively. While 3,722 genes change their expression status (2,854 become active, 868 inactive) during the transition, only 4.2% of genes lose methylation and even fewer genes (0.5%) gain methylation during this transition. Most genes (95%) maintain their methylation status. However, there is a small but significant relationship between the transcriptional change and a change in gene body methylation. Only a very small part of the variation/change is explained by the change in GBM (Fig. 5C). The embryonic gene body methylation level of a gene predicts changes in GBM level between the developmental stages, as higher methylated genes are more likely to be less methylated in the adult stage (Fig. 5B).

### Gene Ontology enrichment indicates functional differences of methylated and not methylated genes

To further characterize which genes are expressed in the four categories, we performed a GO term analysis of the respective subsets of genes for the embryo samples. 258 GO terms were overrepresented compared to the entirety of GO terms associated with all genes in the ‘*methylated / expressed’* group, but only five in the ‘*methylated / not expressed’* group, as this group consists of fewer genes. Among those are transposition related GO terms. ‘*Not methylated / expressed’* genes are frequently assigned to GO terms associated with regulatory functions, while ‘*methylated / expressed’* genes have slightly more GO terms assigned to cellular components (Fig. 6).

**Figure 6.**
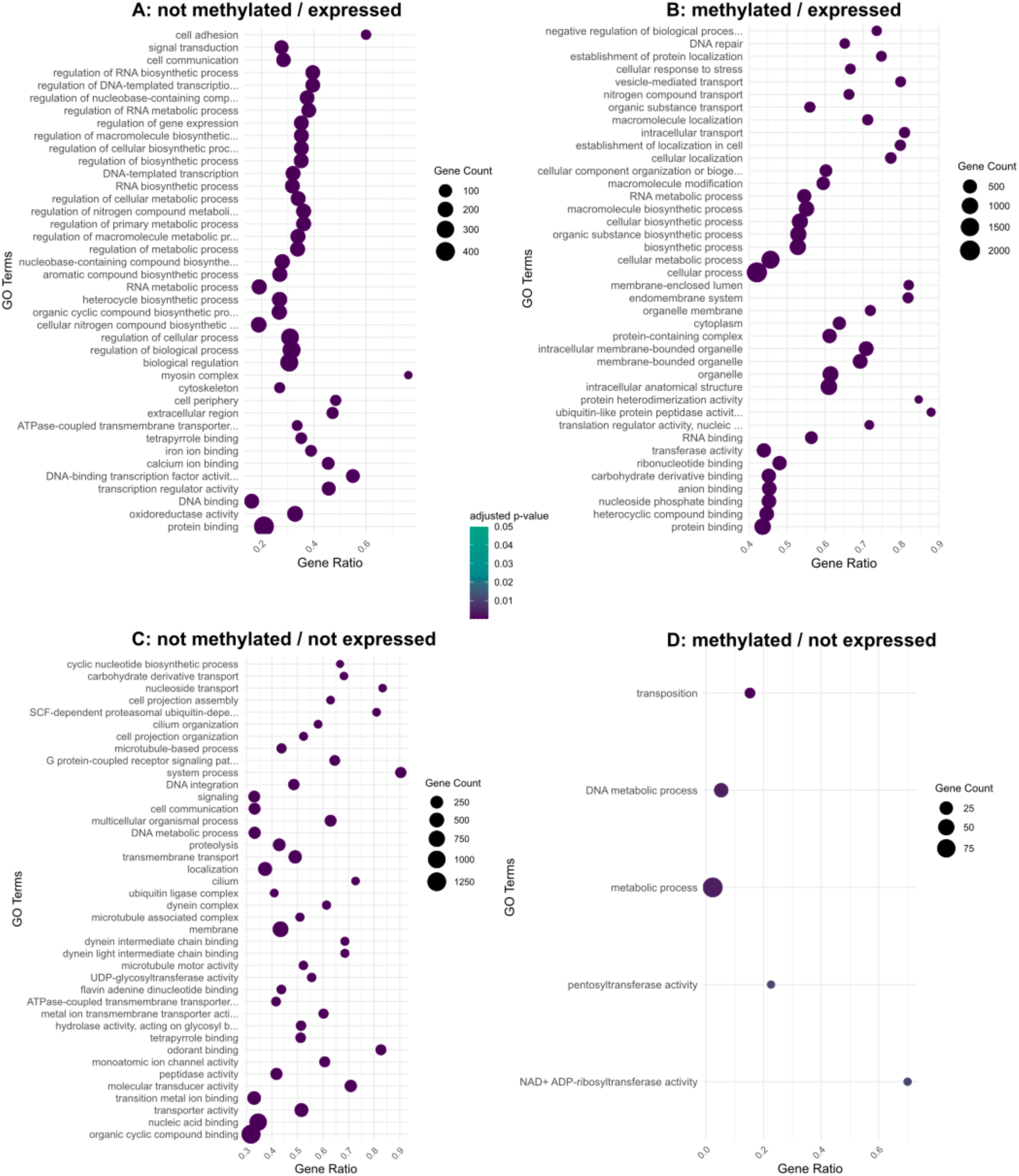
Embryo top 40 significantly enriched Gene Ontology (GO) terms for the four subsets. The p-value was adjusted using the Benjamini-Hochberg procedure. Gene count is the number of genes associated with a GO term, whereas ‘Gene Ratio’ is the percentage of genes of the specific subset in the given GO terms. Categories vary in the number of genes included, resulting in different numbers of enriched GO terms (*‘not methylated / expressed’* = 140 GO; *‘methylated / expressed*’ = 258 GO; *‘not methylated / not expressed*’ = 126 GO; ‘*methylated / not expressed*’ = 5 GO).

*‘Not methylated / expressed’* genes have functional enrichment in DNA binding and transcriptional regulation as key molecular processes. Biological processes include regulation of gene expression and RNA-related heterocycle biosynthetic pathways. ‘*Methylated / expressed’* genes show enrichment for biological processes like intracellular transport, with molecular functions focused on transmembrane transporter activity and RNA binding. Furthermore, methylation-related genes are enriched (39 of 53 entries), chromosome segregation, TOR signaling and RNA modifications. Molecular processes highlight N-acyltransferase activity (20 of 24 entries) and histone-modifying activity (27 of 33 entries). In the most numerous groups of genes, ‘*not methylated / not expressed’*, the enriched GO terms include transporter activity, transmembrane transport, and cellular components like the dynein complex, indicating potentially overarching functions in cellular transport mechanisms.

Due to the high number of shared genes in the respective categories, GO terms for adult categories are overall analogous to embryos and are therefore shown in Fig. S16.

### Only H3K36me3 is associated with CpG methylation, while H3K27ac and H3K36me3 are associated with active transcription

We used CUT&Tag to analyze the enrichment profile of histone modifications H3K27ac and H3K36me3 in embryos. From our initial data set, we selected the 2.5% of highest peaks of each replicate and modification. We used the called peaks from both replicates to create a high confidence set of peaks by taking only the intersections of overlapping peaks into account. Out of this set of most reliable peaks, the majority are covering genes, namely 79% and 83% for H3K27ac and H3K36me3, respectively, with a remaining 21% and 17% being placed in intergenic regions.

Both H3K27ac and H3K36me3 are associated with active gene expression. 66% of ‘*expressed’* genes show an overlap with high confidence peaks for either H3K27ac or H3K36me3. In contrast, this is only the case for 10% of ‘*not expressed’* genes. Furthermore, 58% of expressed genes show enrichment for both modifications simultaneously, which is only the case for 4% of not expressed genes (Fig. S17). A prominent, narrow peak for H3K27ac can be seen at the transcription start site (TSS) of expressed genes (see Fig. 7A). We also observe a small dip of H3K27ac levels around the transcription end site (TES). In inactive genes, only low levels of H3K27ac are observed, with a tiny peak at the TSS followed by a plateau running all the way to the TES (see Fig. 7B). In contrast to the tall and narrow peak of H3K27ac at the TSS, H3K36me3 enrichment presents itself with a steep increase from the TSS towards the TES of the gene, followed by a gradual decline reaching far into the downstream flanking region (see Fig. 7A). The overall expression of ‘*methylated’* genes is higher, with a more pronounced effect in the presence of H3K36me3 or H3K27ac. When both modifications are present, the highest FPKM scores are observed (see Fig.7C-D).

**Figure 7.**
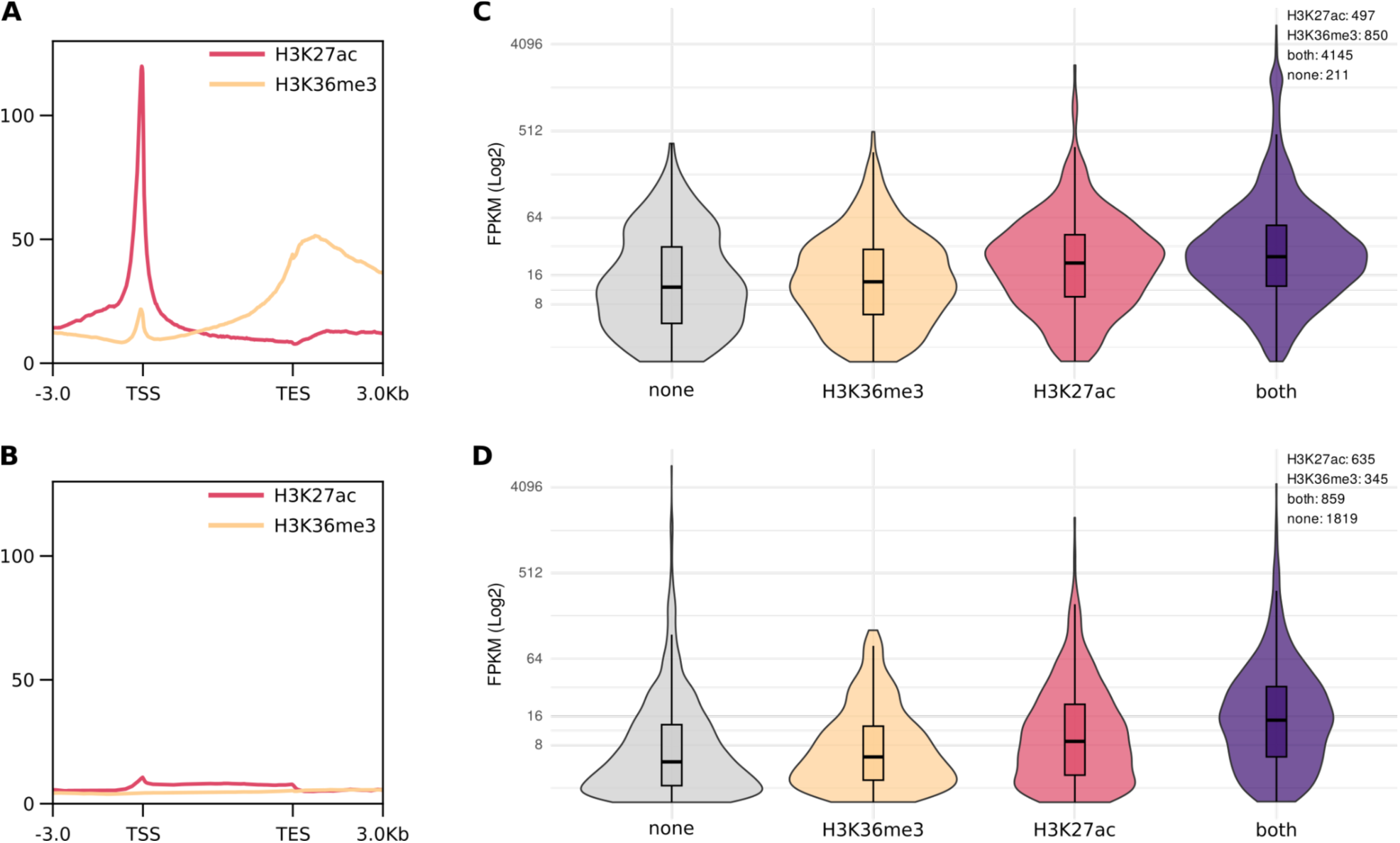
Enrichment patterns of H3K27ac and H3K36me3 in *‘expressed’*. **(A) and *‘not expressed’* (B) genes.** Shown are the profiles for replicate 1 (replicate 2 see Fig. S18). **Differences in expression levels of (C) *‘methylated’* and (D) *‘not methylated’* active genes** in relation to their histone modifications.

Comparing the four categories, as expected, the H3K27ac enrichment is very strong in expressed genes, but does not differ between expressed genes that are ‘*methylated’* or ‘*not methylated’*. This suggests that gene body methylation and H3K27ac are largely independent (see Fig. 8).

**Figure 8.**
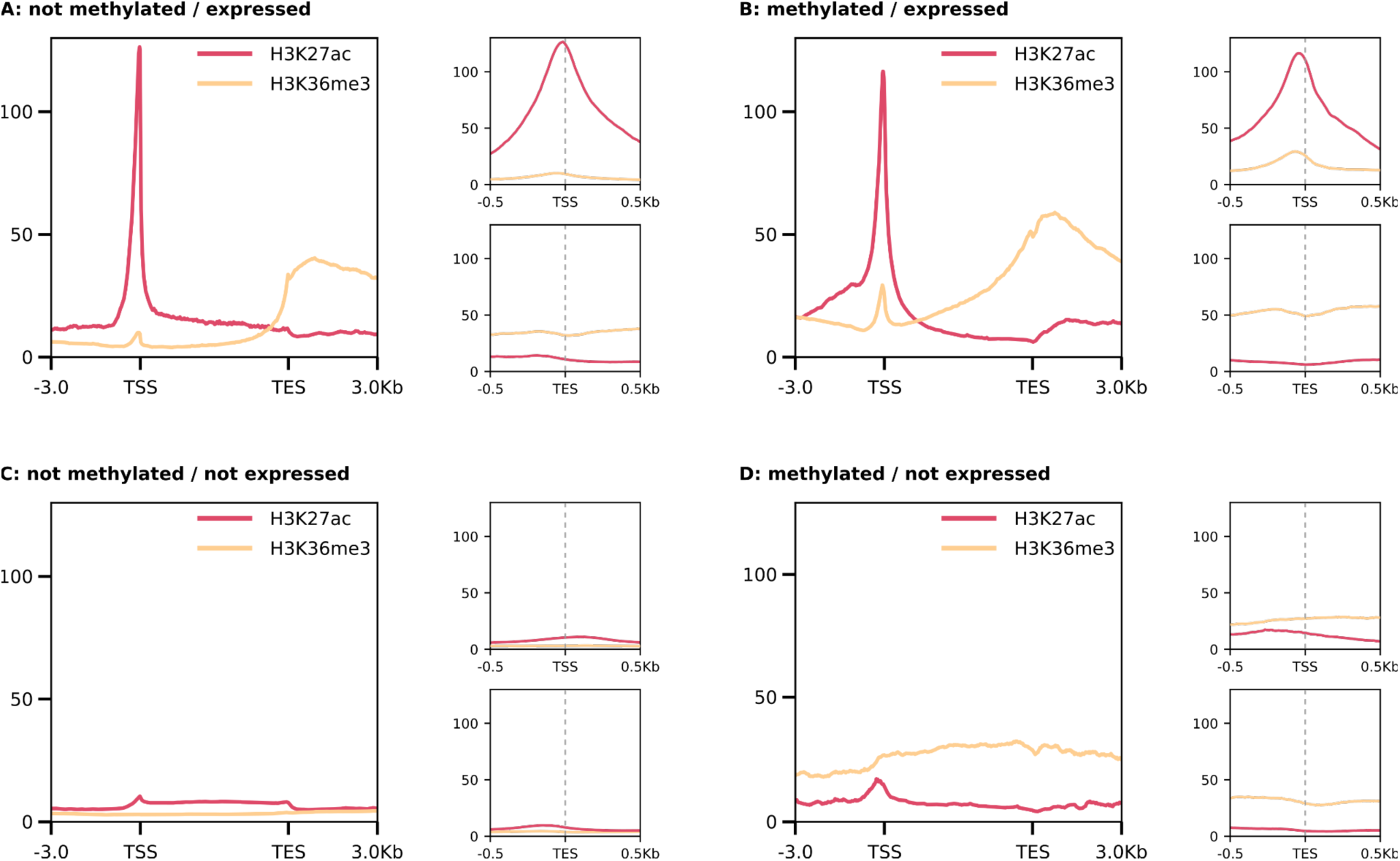
Enrichment profiles of the H3K27ac and H3K36me3 distribution at gene body, transcription start site (TSS) and transcription end site (TES). Genes are normalized to a length of 5 kb, with 3 kb upstream and downstream of TSS and TES, respectively. For TSS and TES, flanking regions of 0.5 kb were used. Shown are the profiles of replicate 1 (profiles for replicate 2 and heatmaps for all replicates see Fig. S19).

H3K36me3 enrichment levels are strongest in the category of ‘*methylated / expressed*’ genes. In contrast to H3K27ac, H3K36me3 profiles show notable differences between ‘*methylated’* and ‘*not methylated’* genes. While *‘methylated’* genes exhibit H3K36me3 enrichment throughout the entire gene body, gradually increasing towards the TES, *not ‘methylated / expressed’* genes show little enrichment throughout the gene body and a steep increase at the end of the gene. Here, the peak at the TSS is visible, but smaller in size. For ‘*methylated / not expressed’* genes we observe a notable level of H3K36me3 enrichment without discernible peaks and the enrichment along the entire gene body is almost equal to the flanking regions. In ‘*not methylated / not expressed’* genes, no enrichment is observed and the small peak of H3K36me3 at the TSS is absent in inactive genes.

## Discussion

In this study, we investigated for the first time the genome-wide epigenetic landscape of a species that has lost the *de novo* DNA methyltransferase DNMT3 but retained the maintenance DNA methyltransferase DNMT1 and CpG methylation, the Colorado potato beetle *L. decemlineata*. Despite the loss of DNMT3, the CpG methylation pattern mirrored H3K36me3 enrichment, with prominent differences in expressed and non-expressed genes, while H3K27ac showed a prominent peak at the transcription start of active genes but did not show any further associations with CpG methylation.

For our multiomics approach in the Colorado potato beetle, a non-model species in molecular biology, we applied new techniques, EM-seq and CUT&Tag, to study CpG methylation and histone modifications, respectively. To our knowledge, it is the first time EM-seq has been applied to an arthropod species. EM-seq promises greater uniformity of GC coverage, and a greater coverage of CpGs at a lower required sequencing coverage depth compared to whole genome bisulfite sequencing (WGBS) (Vaisvila et al., 2021). To generate genome-wide histone modification maps for H3K27ac and H3K36me3, we used CUT&Tag, a more sensitive alternative to ChIP-seq, that requires a lower number of cells, making it particularly suited for small organisms like insects (Kaya-Okur et al., 2019, 2020).

We observed genome-wide CpG methylation levels of 4.2% in embryos and 3.4% in adults, which is rather high compared to other coleopteran species (*T. castaneum*: loss of DNMT3 and CpG methylation (Schulz et al., 2018), *N. vespilloides*: CpG methylation below 1% (Cunningham et al., 2015)) and the previously reported levels in *L. decemlineata* (Brevik et al., 2021). Increased sensitivity of EM-seq compared to WGBS (Vaisvila et al., 2021) or improved genome assembly might contribute to our higher methylation levels.

In invertebrates, unlike vertebrates, DNA methylation does not silence inactive genes (Field et al., 2004); instead, gene body methylation in insects often correlates with active transcription (Zemach et al., 2010). The fraction of methylated genes varies strongly among insect genomes. In *L. decemlineata*, around 25% of annotated genes are methylated to varying degrees, showing a nearly flat distribution between 10-70%, unlike the bimodal pattern seen in Hymenoptera (Dixon & Matz, 2022). While gene body methylation is linked to active transcription, its regulatory role is debated. Studies suggest higher methylation correlates with increased gene expression (Foret et al., 2009; Lewis et al., 2020; Provataris et al., 2018), and we observed a positive, though small correlation, supporting the idea that gene body methylation facilitates smooth transcription rather than regulating it (de Mendoza et al., 2020; Dixon & Matz, 2022).

In *L. decemlineata*, exons are more highly methylated than introns or flanking regions, with increased methylation in later exons of expressed genes, suggesting potential regulatory relevance. This aligns with an unclear trend in invertebrates, where preferentially methylated exons vary: *N. vespilloides* shows higher methylation in the first three exons, while *Blattella germanica* has elevated methylation starting from exon four (Lewis et al., 2020). Methylation drops at transcription start sites (TSS) in expressed genes, a feature absent in not expressed genes. Unmethylated promoters are essential for transcriptional initiation in vertebrates (Isagawa et al., 2011). While unmethylated promoters are common in insects, some exceptions (e.g., *Planococcus citri* and *Strigamia maritima*) show methylated promoters (Lewis et al., 2020). Reduced TSS methylation, as previously reported in the purple sea urchin *Strongylocentrotus purpuratus*, may enhance chromatin accessibility and gene expression also in our study organism (Bogan et al., 2023).

In *L. decemlineata* ‘*methylated / expressed’* genes were, on average, the longest, which is similar to other invertebrates but contrasting to honeybee and silk moth, where highly methylated genes are shorter compared to lowly methylated genes (Sarda et al., 2012). However, these associations could be species specific and even without further regulatory relevance.

Using GO analysis, we found that *‘Not methylated / expressed’* genes are often linked to regulatory processes requiring flexible expression, while ‘*methylated / expressed’* genes are associated with stable expression and roles in DNA repair and stress responses. Methylation in active genes may support stable regulation, whereas unmethylated genes allow for more dynamic expression, mediated by other transcriptional regulators. Consistent with this, we observed that not methylated genes more frequently switch expression between embryo and adult stages compared to methylated genes. ‘*Methylated / not expressed’* genes lack a TSS dip, suggesting that methylation may suppress transcription by restricting transcription factor access, likely reducing expression of potentially harmful genes, such as those with transposase activity.

As the overarching goal of our study was to uncover how the loss of DNMT3 affects the methylation landscape and the distribution of histone modification, we studied embryonic H3K36me3 and H3K27ac patterns. In vertebrates, both histone modifications are involved in marking active genes. The interaction between H3K36me3 and methylated cytosine is deeply conserved in evolution. For this reason, H3K36me3 patterns in *D. melanogaster* (lacking a CpG methylation system) can predict the CpG methylation landscape in *S. invicta*, *A. mellifera,* and *B. mori* (Hunt et al., 2013; Nanty et al., 2011). We also find a positive association between the abundance of H3K36me3 and CpG methylation at actively expressed genes.

In vertebrates, H3K36me3 enrichment increases towards the 3’-end of a transcribed gene and decays after the TES (Neri et al., 2017). While this is generally reflected in our data, we surprisingly observe high H3K36me3 enrichment levels extending more than 3kb into the downstream flanking region of the gene. Our data also corroborates the enrichment of H3K36me3 at the TSS, as previously described (Zhang et al., 2022), though the level of H3K36me3 is significantly lower at the TSS compared to the TES. This peak may actually be independent of the main, genic H3K36me3 enrichment pattern, since a different enzyme, i.e SMYD5, is possibly responsible for setting this mark (Zhang et al., 2022). In contrast to not methylated and inactive genes, which show no enrichment of H3K36me3, ‘*methylated / not expressed’* display uniform H3K36me3 enrichment across the entire gene body. In general, the shapes of H3K36me3 enrichment profiles and CpG methylation levels seem to mirror each other.

If DNMT3 is the only enzyme able to methylate regions *de novo* marked by H3K36me3, through recognition by its PWWP domain, then methylation patterns observed from embryo to adult stages in *L. decemlineata* would reflect germline patterns maintained by DNMT1, with some methylation loss in somatic cells. This aligns with our findings of reduced methylation from embryonic to adult stages, though some genes appear to gain methylation during development. Further studies could explore if and how CpG methylation is acquired in *L. decemlineata*.

H3K27ac, unlike the neutral methylation marks, directly increases DNA accessibility and likely marked active genes in early eukaryotes (Prohaska et al., 2010; Iyer et al., 2008). In vertebrates, H3K27ac is found on active genes and enhancers, and our data shows a prominent TSS peak indicating active transcription. While enzymes like CBP/p300 add H3K27ac, (Xu et al., 2021)) suggest MBD2/3 binds to intragenic methylated CpG near the TSS, recruiting Tip60 acetyltransferase to promote H3K27ac. However, we observed that H3K27ac is tied to transcriptional status rather than CpG methylation, challenging the generality of the previously reported link between H3K27ac and CpG methylation (Xu et al., 2021).

Histone modifications regulate transcriptional processes in a concerted fashion. Several signals need to come together before transcription can be initiated. Histone marks “remember the past and predict the future”, meaning that parts of a composite pattern are preset, building up to a future exertion. Likewise, the deconstruction of a composite epigenetic pattern may be incomplete leaving traces of past actions. This might explain why we observe variance in associations of histone modifications with DNA methylation and gene expression.

## Conclusion

We studied the genome-wide CpG distribution in *L. decemlineata* using EM-seq for the first time in an insect species. While levels of DNA methylation were surprisingly high compared to other coleopteran species, we found as expected exons to be most highly enriched in CpG methylation. Consistent with the loss of DNMT3 in this species, we observed a reduction of methylation in adults compared to embryos. On the other hand, the loss of DNMT3 did not seem to affect the association between CpG methylation and H3K36me3 enrichment that are mirroring each other. This similarity consistently remains with differing patterns in *‘methylated / expressed’* vs. *‘methylated / not expressed’* genes. The H3K36me3 enrichment extends more than 3kb into the downstream flanking regions of genes. H3K27ac apparently does not seem to have an association with CpG methylation, while peaks of H3K27ac predict gene expression. Taken together, our study demonstrates that connections between the different epigenetic systems show evolutionary flexibility and therefore urges caution regarding too simplistic generalizations.

## Author Contributions

S.J.P and J.K. designed and supervised the study. Z.M.L. conducted the laboratory experiment. Z.M.L, A.K. and C.I.K.V. did the CUT&Tag experiment. Z.M.L., E.I., and J.E. analyzed the data. Z.M.L. and E.I. wrote the manuscript. S.J.P and J.K. revised the manuscript. All authors read, commented and approved the final version of the manuscript.

## Acknowledgements

We would like to thank Prof. Shuqing Xu (University of Mainz) for kindly providing us with the initial Colorado potato beetle population. We thank the IMB Genomics core facility for their support. We furthermore want to thank the Colone Center for Genomics for their support and expertise with the Enzymatic methyl sequencing. This study was funded by the Deutsche Forschungsgemeinschaft (DFG, German Research Foundation) – priority program SPP 2349, project number 503 349 225 to S.J.P. and J.K.

## Data availability statement

Additional data supporting this article are included in the supplementary file.

## Literature

Adrian Alexa, J. R. (2017). topGO. Bioconductor. 10.18129/B9.BIOC.TOPGO

Aliaga, B., Bulla, I., Mouahid, G., Duval, D., & Grunau, C. (2019). Universality of the DNA methylation codes in Eucaryotes. Scientific Reports, 9(1), 1–11.

Attali, D., & Baker, C. (2015). GgExtra: Add marginal histograms to “ggplot2”, and more “ggplot2” enhancements [Dataset]. In CRAN: Contributed Packages. The R Foundation. 10.32614/cran.package.ggextra

Bogan, S. N., Strader, M. E., & Hofmann, G. E. (2023). Associations between DNA methylation and gene regulation depend on chromatin accessibility during transgenerational plasticity. BMC Biology, 21(1), 1–18.

Bourc’his, D., & Bestor, T. H. (2004). Meiotic catastrophe and retrotransposon reactivation in male germ cells lacking Dnmt3L. Nature, 431(7004), 96–99.

Brevik, K., Bueno, E. M., McKay, S., Schoville, S. D., & Chen, Y. H. (2021). Insecticide exposure affects intergenerational patterns of DNA methylation in the Colorado potato beetle,. Evolutionary Applications, 14(3), 746–757.

Brunson, J. C., & Read, Q. D. (2017). ggalluvial: Alluvial Plots in “ggplot2” [Dataset]. In CRAN: Contributed Packages. The R Foundation. 10.32614/cran.package.ggalluvial

Cardoso-Júnior, C. A. M., Yagound, B., Ronai, I., Remnant, E. J., Hartfelder, K., & Oldroyd, B. P. (2021). DNA methylation is not a driver of gene expression reprogramming in young honey bee workers. Molecular Ecology, 30(19), 4804–4818.

Chen, H. (2011). VennDiagram: Generate high-resolution Venn and Euler plots [Dataset]. In CRAN: Contributed Packages. The R Foundation. 10.32614/cran.package.venndiagram

Cunningham, C. B., Ji, L., Wiberg, R. A. W., Shelton, J., McKinney, E. C., Parker, D. J., Meagher, R. B., Benowitz, K. M., Roy-Zokan, E. M., Ritchie, M. G., Brown, S. J., Schmitz, R. J., & Moore, A. J. (2015). The Genome and Methylome of a Beetle with Complex Social Behavior, Nicrophorus vespilloides (Coleoptera: Silphidae). Genome Biology and Evolution, 7(12), 3383–3396.

de Mendoza, A., Lister, R., & Bogdanovic, O. (2020). Evolution of DNA Methylome Diversity in Eukaryotes. Journal of Molecular Biology, 432(6), 1687–1705.

Dhayalan, A., Rajavelu, A., Rathert, P., Tamas, R., Jurkowska, R. Z., Ragozin, S., & Jeltsch, A. (2010). The Dnmt3a PWWP domain reads histone 3 lysine 36 trimethylation and guides DNA methylation. The Journal of Biological Chemistry, 285(34), 26114–26120.

Dixon, G., & Matz, M. (2022). Changes in gene body methylation do not correlate with changes in gene expression in Anthozoa or Hexapoda. BMC Genomics, 23(1), 1–11.

Edmunds, J. W., Mahadevan, L. C., & Clayton, A. L. (2007). Dynamic histone H3 methylation during gene induction: HYPB/Setd2 mediates all H3K36 trimethylation. The EMBO Journal. 10.1038/sj.emboj.7601967

Engelhardt, J., Scheer, O., Stadler, P. F., & Prohaska, S. J. (2022). Evolution of DNA Methylation Across Ecdysozoa. Journal of Molecular Evolution, 90(1), 56–72.

Field, L. M., Lyko, F., Mandrioli, M., & Prantera, G. (2004). DNA methylation in insects. Insect Molecular Biology, 13(2), 109–115.

Flynn, J. M., Hubley, R., Goubert, C., Rosen, J., Clark, A. G., Feschotte, C., & Smit, A. F. (2020). RepeatModeler2 for automated genomic discovery of transposable element families. Proceedings of the National Academy of Sciences, 117(17), 9451–9457.

Foret, S., Kucharski, R., Pittelkow, Y., Lockett, G. A., & Maleszka, R. (2009). Epigenetic regulation of the honey bee transcriptome: unravelling the nature of methylated genes. BMC Genomics, 10(1), 1–11.

Grau-Bové, X., Navarrete, C., Chiva, C., Pribasnig, T., Antó, M., Torruella, G., Galindo, L. J., Lang, B. F., Moreira, D., López-Garcia, P., Ruiz-Trillo, I., Schleper, C., Sabidó, E., & Sebé-Pedrós, A. (2022). A phylogenetic and proteomic reconstruction of eukaryotic chromatin evolution. Nature Ecology & Evolution, 6(7), 1007–1023.

Greenberg, M. V. C., & Bourc’his, D. (2019). The diverse roles of DNA methylation in mammalian development and disease. Nature Reviews. Molecular Cell Biology, 20(10), 590–607.

Guenther, M. G., Levine, S. S., Boyer, L. A., Jaenisch, R., & Young, R. A. (2007). A chromatin landmark and transcription initiation at most promoters in human cells. Cell, 130(1), 77–88.

Hahn, M. A., Wu, X., Li, A. X., Hahn, T., & Pfeifer, G. P. (2011). Relationship between Gene Body DNA Methylation and Intragenic H3K9me3 and H3K36me3 Chromatin Marks. PloS One, 6(4), e18844.

Harrell, F. E., Jr. (2003). Hmisc: Harrell Miscellaneous [Dataset]. In CRAN: Contributed Packages. The R Foundation. 10.32614/cran.package.hmisc

Harris, K. D., Lloyd, J. P. B., Domb, K., Zilberman, D., & Zemach, A. (2019). DNA methylation is maintained with high fidelity in the honey bee germline and exhibits global non-functional fluctuations during somatic development. Epigenetics & Chromatin, 12(1), 1–18.

Hunt, B. G., Glastad, K. M., Yi, S. V., & Goodisman, M. A. D. (2013). The function of intragenic DNA methylation: insights from insect epigenomes. Integrative and Comparative Biology, 53(2), 319– 328.

Isagawa, T., Nagae, G., Shiraki, N., Fujita, T., Sato, N., Ishikawa, S., Kume, S., & Aburatani, H. (2011). DNA Methylation Profiling of Embryonic Stem Cell Differentiation into the Three Germ Layers. PloS One, 6(10), e26052.

Iyer, L. M., Anantharaman, V., Wolf, M. Y., L. Aravind, L. (2008). Comparative genomics of transcription factors and chromatin proteins in parasitic protists and other eukaryotes. International Journal for Parasitology, 38(1), 1–31.

Jühling, F., Kretzmer, H., Bernhart, S. H., Otto, C., Stadler, P. F., & Hoffmann, S. (2016). metilene: fast and sensitive calling of differentially methylated regions from bisulfite sequencing data. Genome Research, 26(2), 256–262.

Kang, Y., Kim, Y. W., Kang, J., & Kim, A. (2021). Histone H3K4me1 and H3K27ac play roles in nucleosome eviction and eRNA transcription, respectively, at enhancers. FASEB Journal: Official Publication of the Federation of American Societies for Experimental Biology, 35(8), e21781.

Kaya-Okur, H. S., Janssens, D. H., Henikoff, J. G., Ahmad, K., & Henikoff, S. (2020). Efficient low-cost chromatin profiling with CUT&Tag. Nature Protocols, 15(10), 3264–3283.

Kaya-Okur, H. S., Wu, S. J., Codomo, C. A., Pledger, E. S., Bryson, T. D., Henikoff, J. G., Ahmad, K., & Henikoff, S. (2019). CUT&Tag for efficient epigenomic profiling of small samples and single cells. Nature Communications, 10(1), 1–10.

Krueger, F., & Andrews, S. R. (2011). Bismark: a flexible aligner and methylation caller for Bisulfite-Seq applications. Bioinformatics, 27(11), 1571–1572.

Krueger, F., James, F., Ewels, P., Afyounian, E., Weinstein, M., Schuster-Boeckler, B., Hulselmans, G., & sclamons. (2023). FelixKrueger/TrimGalore: v0.6.10 - add default decompression path. Zenodo. 10.5281/ZENODO.7598955

Langmead, B., & Salzberg, S. L. (2012). Fast gapped-read alignment with Bowtie 2. Nature Methods, 9(4), 357–359.

Lewis, S. H., Ross, L., Bain, S. A., Pahita, E., Smith, S. A., Cordaux, R., Miska, E. A., Lenhard, B., Jiggins, F. M., & Sarkies, P. (2020) Widespread conservation and lineage-specific diversification of genome-wide DNA methylation patterns across arthropods. PLoS Genetics, 16(6), e1008864.

Lister, R., Pelizzola, M., Dowen, R. H., Hawkins, R. D., Hon, G., Tonti-Filippini, J., Nery, J. R., Lee, L., Ye, Z., Ngo, Q.-M., Edsall, L., Antosiewicz-Bourget, J., Stewart, R., Ruotti, V., Millar, A. H., Thomson, J. A., Ren, B., & Ecker, J. R. (2009). Human DNA methylomes at base resolution show widespread epigenomic differences. Nature, 462(7271), 315–322.

Love, M. I., Huber, W., & Anders, S. (2014). Moderated estimation of fold change and dispersion for RNA-seq data with DESeq2. Genome Biology, 15(12), 550.

Lyko, F. (2017). The DNA methyltransferase family: a versatile toolkit for epigenetic regulation. Nature Reviews. Genetics, 19(2), 81–92.

Maleszka, R. (2024). Reminiscences on the honeybee genome project and the rise of epigenetic concepts in insect science. Insect Molecular Biology, 33(5), 444–456.

Martin, M. (2011). Cutadapt removes adapter sequences from high-throughput sequencing reads. EMBnet.journal, 17(1), 10–12.

Meers, M. P., Tenenbaum, D., & Henikoff, S. (2019). Peak calling by Sparse Enrichment Analysis for CUT&RUN chromatin profiling. Epigenetics & Chromatin, 12(1), 1–11.

Mortazavi, A., Williams, B. A., McCue, K., Schaeffer, L., & Wold, B. (2008). Mapping and quantifying mammalian transcriptomes by RNA-Seq. Nature Methods, 5(7), 621–628.

Nanty, L., Carbajosa, G., Heap, G. A., Ratnieks, F., van Heel, D. A., Down, T. A., & Rakyan, V. K. (2011). Comparative methylomics reveals gene-body H3K36me3 in Drosophila predicts DNA methylation and CpG landscapes in other invertebrates. Genome Research, 21(11), 1841–1850.

Nègre, N., Brown, C. D., Ma, L., Bristow, C. A., Miller, S. W., Wagner, U., Kheradpour, P., Eaton, M. L., Loriaux, P., Sealfon, R., Li, Z., Ishii, H., Spokony, R. F., Chen, J., Hwang, L., Cheng, C., Auburn, R. P., Davis, M. B., Domanus, M., … White, K. P. (2011). A cis-regulatory map of the Drosophila genome. Nature, 471(7339), 527–531.

Neri, F., Rapelli, S., Krepelova, A., Incarnato, D., Parlato, C., Basile, G., Maldotti, M., Anselmi, F., & Oliviero, S. (2017). Intragenic DNA methylation prevents spurious transcription initiation. Nature, 543(7643), 72–77.

Pokholok, D. K., Harbison, C. T., Levine, S., Cole, M., Hannett, N. M., Lee, T. I., Bell, G. W., Walker, K., Rolfe, P. A., Herbolsheimer, E., Zeitlinger, J., Lewitter, F., Gifford, D. K., & Young, R. A. (2005). Genome-wide map of nucleosome acetylation and methylation in yeast. Cell, 122(4), 517–527.

Prohaska, S. J., Stadler, P. F., Krakauer, D. C. (2010). Innovation in gene regulation: The case of chromatin computation. Journal of Theoretical Biology, 265(1), 27–44.

Provataris, P., Meusemann, K., Niehuis, O., Grath, S., & Misof, B. (2018). Signatures of DNA Methylation across Insects Suggest Reduced DNA Methylation Levels in Holometabola. Genome Biology and Evolution, 10(4), 1185–1197.

Quinlan, A. R., & Hall, I. M. (2010). BEDTools: a flexible suite of utilities for comparing genomic features. Bioinformatics, 26(6), 841–842.

Raddatz, G., Guzzardo, P. M., Olova, N., Fantappié, M. R., Rampp, M., Schaefer, M., Reik, W., Hannon, G. J., & Lyko, F. (2013). Dnmt2-dependent methylomes lack defined DNA methylation patterns. Proceedings of the National Academy of Sciences, 110(21), 8627–8631.

Ramírez, F., Ryan, D. P., Grüning, B., Bhardwaj, V., Kilpert, F., Richter, A. S., Heyne, S., Dündar, F., & Manke, T. (2016). deepTools2: a next generation web server for deep-sequencing data analysis. Nucleic Acids Research, 44(W1), W160–W165.

Sarda, S., Zeng, J., Hunt, B. G., & Yi, S. V. (2012). The Evolution of Invertebrate Gene Body Methylation. Molecular Biology and Evolution, 29(8), 1907–1916.

Schulz, N. K. E., Wagner, C. I., Ebeling, J., Raddatz, G., Diddens-de Buhr, M. F., Lyko, F., & Kurtz, J. (2018). Dnmt1 has an essential function despite the absence of CpG DNA methylation in the red flour beetle Tribolium castaneum. Scientific Reports, 8(1), 16462.

Simola, D. F., Wissler, L., Donahue, G., Waterhouse, R. M., Helmkampf, M., Roux, J., Nygaard, S., Glastad, K. M., Hagen, D. E., Viljakainen, L., Reese, J. T., Hunt, B. G., Graur, D., Elhaik, E., Kriventseva, E. V., Wen, J., Parker, B. J., Cash, E., Privman, E., … Gadau, J. (2013). Social insect genomes exhibit dramatic evolution in gene composition and regulation while preserving regulatory features linked to sociality. Genome Research, 23(8), 1235–1247.

Suzuki, M. M., Kerr, A. R. W., De Sousa, D., & Bird, A. (2007). CpG methylation is targeted to transcription units in an invertebrate genome. Genome Research, 17(5), 625–631.

Teissandier, A., & Bourc’his, D. (2017). Gene body DNA methylation conspires with H3K36me3 to preclude aberrant transcription. The EMBO Journal, 36(11), 1471–1473.

Vaisvila, R., Ponnaluri, V. K. C., Sun, Z., Langhorst, B. W., Saleh, L., Guan, S., Dai, N., Campbell, M. A., Sexton, B. S., Marks, K., Samaranayake, M., Samuelson, J. C., Church, H. E., Tamanaha, E., Corrêa, I. R., Jr, Pradhan, S., Dimalanta, E. T., Evans, T. C., Jr, Williams, L., & Davis, T. B. (2021). Enzymatic methyl sequencing detects DNA methylation at single-base resolution from picograms of DNA. Genome Research, 31(7), 1280–1289.

Wagner, E. J., & Carpenter, P. B. (2012). Understanding the language of Lys36 methylation at histone H3. Nature Reviews. Molecular Cell Biology, 13(2), 115–126.

Wickham, H. (2007). Reshaping Data with thereshapePackage. Journal of Statistical Software, 21(12). 10.18637/jss.v021.i12

Wickham, H., Chang, W., Henry, L., Pedersen, T. L., Takahashi, K., Wilke, C., Woo, K., Yutani, H., Dunnington, D., & van den Brand, T. (2007). Ggplot2: Create elegant data visualisations using the grammar of graphics [Dataset]. In CRAN: Contributed Packages. The R Foundation. 10.32614/cran.package.ggplot2

Wickham, H., François, R., Henry, L., Müller, K., & Vaughan, D. (2014). dplyr: A Grammar of Data Manipulation [Dataset]. In CRAN: Contributed Packages. The R Foundation. 10.32614/cran.package.dplyr

Wickham, H., Vaughan, D., & Girlich, M. (2014). tidyr: Tidy Messy Data [Dataset]. In CRAN: Contributed Packages. The R Foundation. 10.32614/cran.package.tidyr

Wilhelm, L., Wang, Y., & Xu, S. (2024). The Colorado potato beetle gene expression atlas. In bioRxiv (p. 2024.03.28.587222). 10.1101/2024.03.28.587222

Xu, G., Lyu, H., Yi, Y., Peng, Y., Feng, Q., Song, Q., Gong, C., Peng, X., Palli, S. R., & Zheng, S. (2021). Intragenic DNA methylation regulates insect gene expression and reproduction through the MBD/Tip60 complex. iScience, 24(2), 102040.

Yan, J., Zhang, C., Zhang, M., Zhou, H., Zuo, Z., Ding, X., Zhang, R., Li, F., & Gao, Y. (2023). Chromosome-level genome assembly of the Colorado potato beetle, Leptinotarsa decemlineata. Scientific Data, 10(1), 1–7.

Yano, S., Ishiuchi, T., Abe, S., Namekawa, S. H., Huang, G., Ogawa, Y., & Sasaki, H. (2022). Histone H3K36me2 and H3K36me3 form a chromatin platform essential for DNMT3A-dependent DNA methylation in mouse oocytes. Nature Communications, 13(1), 4440.

Yoh, S. M., Lucas, J. S., & Jones, K. A. (2008). The Iws1:Spt6:CTD complex controls cotranscriptional mRNA biosynthesis and HYPB/Setd2-mediated histone H3K36 methylation. Genes & Development, 22(24), 3422–3434.

Zemach, A., McDaniel, I. E., Silva, P., & Zilberman, D. (2010). Genome-wide evolutionary analysis of eukaryotic DNA methylation. Science, 328(5980), 916–919.

Zhang, Y., Fang, Y., Tang, Y., Han, S., Jia, J., Wan, X., Chen, J., Yuan, Y., Zhao, B., & Fang, D. (2022). SMYD5 catalyzes histone H3 lysine 36 trimethylation at promoters. Nature Communications, 13(1), 1–19.

