## Supplementary Material for "Multiomics reveal associations between CpG methylation, histone modifications and transcription in a species that has lost DNMT3, the Colorado potato beetle"

### Supplementary methods

#### Model organism and samples

We reared Colorado potato beetles, *Leptinotarsa decemlineata* (Coleoptera), in non-overlapping generations on approximately 6-week-old potato (*Solanum tuberosum)* plants (Annabelle variety, purchased from Ellenberg´s Kartoffelvielfalt GmbH & Co. KG, Barum, Germany) in mesh cages (47.5 x 47.5 x 47.5 cm, BugDorm-4E4545, Megaview, Taichung, Taiwan) under constant conditions (16:8 light dark cycle, 70% humidity, 24 °C). Plants were grown in plastic trays in 'Topfsubstrat D400', Stender, in a greenhouse (16:8 light dark cycle) at 24 °C. All life stages were allowed to feed on the plant freely.

#### DNA methylation: Enzymatic methyl sequencing (EM-seq)

We submerged flash-frozen *L. decemlineata* (embryo/adult) in 195 µL TNES buffer (400mM NaCl, 200mM EDTA, 50 mM Tris pH 8.0, 0.5 % SDS) with 5 µL proteinase K (20 mg/mL) and homogenized with a sterile pestle. The homogenate was incubated at 55°C for 1h. After centrifugation (18000 g, 5min), the supernatant was taken, and an equal volume of 24:1 chloroform:isoamyl was added. We mixed by inversion and centrifuged at 10000 x g for 2 min. The aqueous phase was transferred and 60 µL NaCl and 290 µL 96 % Ethanol were added and mixed by inversion. The solution was then stored at -20 °C for 1 h and centrifuged (18000 x g, 15min) afterwards. We removed the supernatant and added 1 mL 70 % chilled Ethanol before centrifuging at 18000 x g for 5 min. Washing was repeated once, then the supernatant was removed and the pellet was left for air dry. 40 µL elution buffer (Tris-HCl pH 8.5) was added. To resuspend the pellet, we heated it to 50 °C for 3min. 2µL RNAse A was added and it was incubated at 37 °C for 30 min. DNA was cleaned before library preparation with the DNeasy PowerClean Kit, Qiagen, following the manufacturer's instructions.

**Table S1. Enzymatic methyl sequencing.** The number of reads, percentage of uniquely mapped reads and coverage of the 3 embryo and 3 adult *L. decemlineata* replicates.

| | **reads** | **% uniquely mapped (average after deduplication)** | **coverage** |
|---|---|---|---|
| **Adult 1** | 64 mio | 50.2% | 8.7X |
| **Adult 2** | 61 mio | | |
| **Adult 3** | 81 mio | | |
| **Embryo 1** | 88 mio | 42.4% | 8.3X |
| **Embryo 2** | 81 mio | | |
| **Embryo 3** | 60 mio | | |

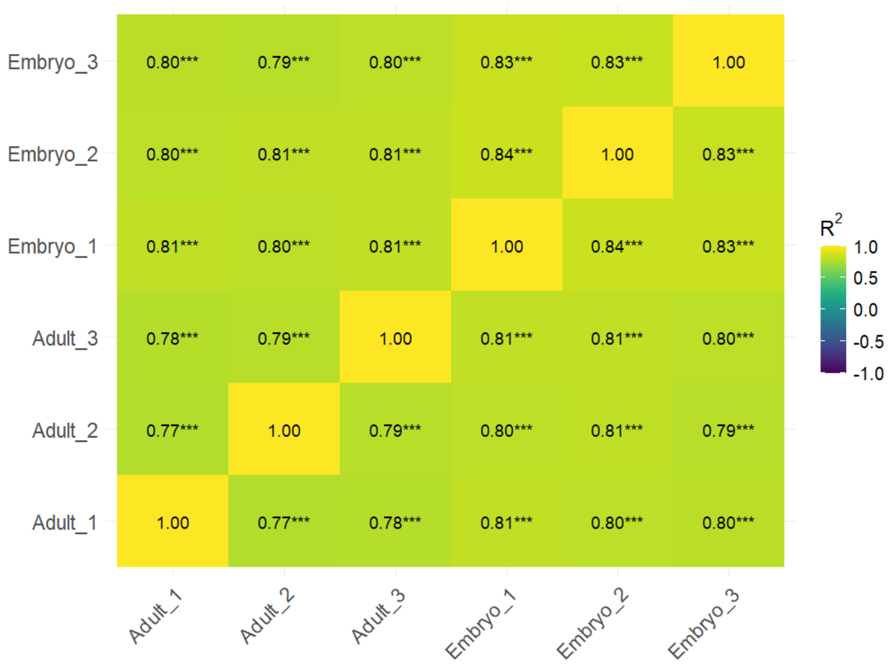


**Figure S1. Correlation matrix showing the Pearson correlation coefficients between biological replicates and life stages of *L. decemlineata* based on genomic CpG sites.** Pearson correlation coefficients (R2) were calculated for the methylation levels per CpG site. Significant correlations ($p < 0.05$) are marked with stars. The correlation strength is color coded according to the legend, with yellow and green shades indicating stronger positive correlations.

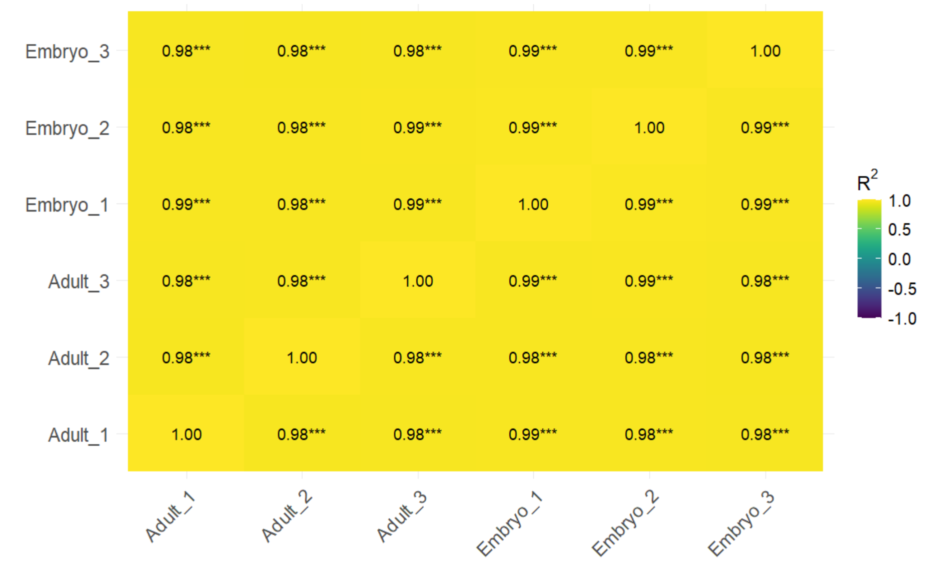


**Figure S2. Correlation matrix showing the Pearson correlation coefficients between biological replicates and life stages of *L. decemlineata* based on genes.** Pearson correlation coefficients (R2) were calculated for the mean % CpG methylation per gene. Significant correlations ($p < 0.05$) are marked with stars. The correlation strength is color coded according to the legend, with yellow and green shades indicating stronger positive correlations.

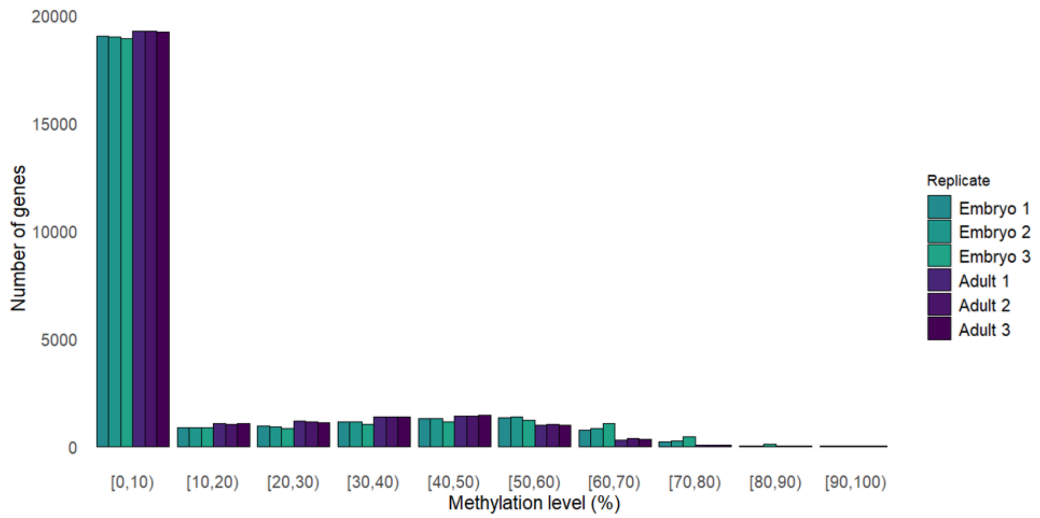


**Figure S3. Number of genes in each methylation percentage range for each replicate of embryo and adult *L. decemlineata*.**

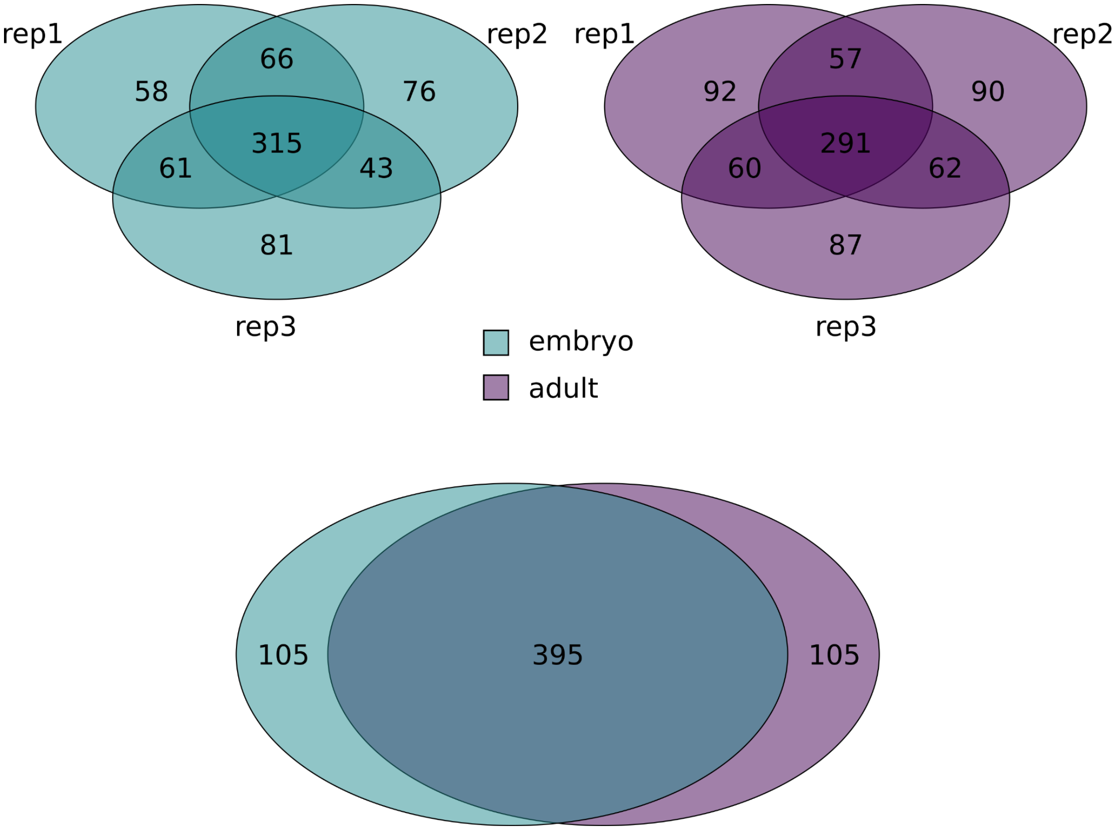


**Figure S4. Overlap of the 500 genes with the highest gene body methylation (%) in *L. decemlineata* between A) the embryonic replicates.** One embryonic replicate consists of 30 pooled embryos of the same age (+- 1h). **B) Between adult replicates and C) between embryos and adults.** The mean methylation level of the 3 replicates of each life stage was used.

**Gene expression: RNA extraction and RNA-seq**

We used a protocol combining Trizol lysis and chloroform extraction with the purification via spin columns from the SV Total RNA Isolation System (Promega). Here, 500 µl of Trizol was added to the frozen sample, which was then homogenized with a clean, sterile pestle. Another 500 µl of Trizol was added to the homogenized sample. The samples were sonicated for 10 minutes and incubated for another 10 minutes while being regularly vortexed. After a 5 min centrifugation (13000 rpm, 4 °C), the supernatant was transferred to a new 1.5 ml Eppendorf tube. 200 µl chloroform was added to the supernatant and the mix was incubated for 15 minutes under regular inversion and centrifuged afterward at 10500 rcf, 15 min, 4 °C). Following the manual instruction, The liquid phase was taken to continue with the purification via spin column.

**Table S2. Number of reads and percentage of uniquely mapped reads of 3 adult and the two remaining embryo *L. decemlineata* RNA-seq replicates.**

| | **reads** | **% uniquely mapped** |
|---|---|---|
| **adult 1** | 62 mio | 76% |
| **adult 2** | 75 mio | 60% |
| **adult 3** | 68 mio | 70% |
| **embryo 1** | removed | removed |
| **embryo 2** | 64 mio | 75% |
| **embryo 3** | 69 mio | 78% |

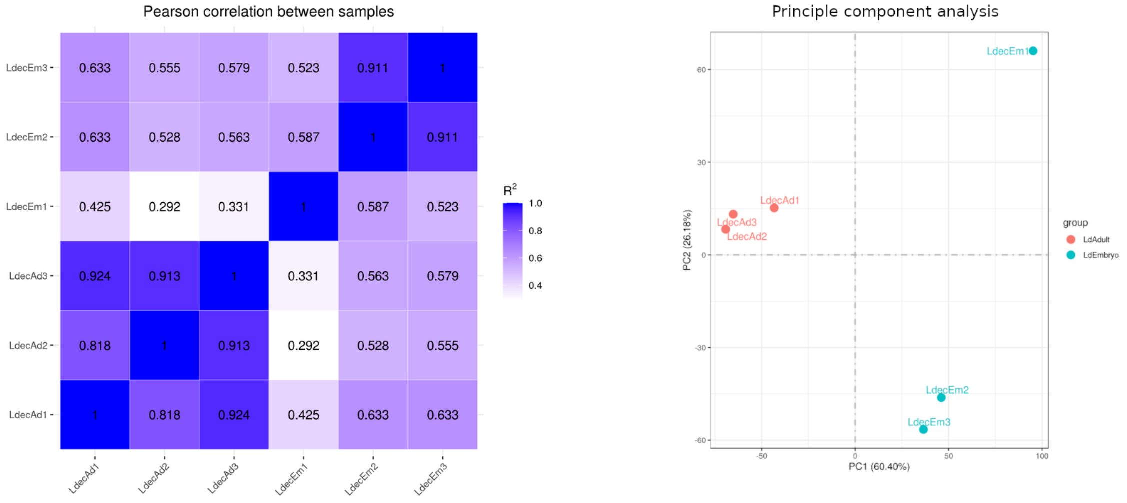


**Figure S5. Pearson correlation and Principal Component Analysis of the 3 embryonic and 3 adult *L. decemlineata* RNA-seq replicates.**

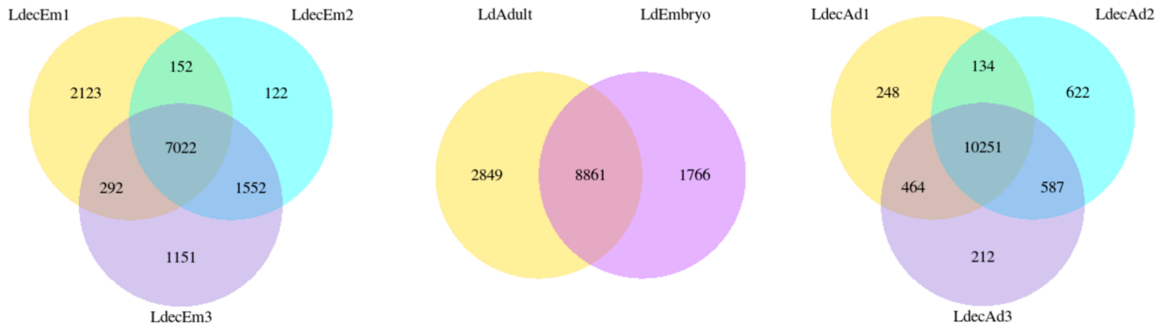


**Figure S6. RNA-seq results as obtained by Novogene.** The embryonic replicate LdecE1 does not correlate or cluster with the other embryonic or adult replicates and was therefore removed from further analysis.

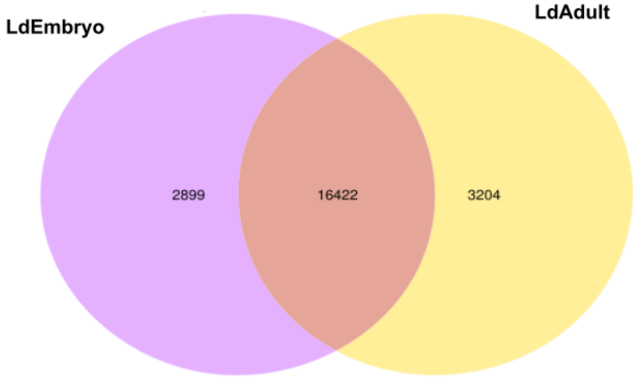


**Figure S7. Comparison of LdAdult and LdEmbryo after removal of embryo 1.**

**Histone modifications: CUT&Tag library generation and sequencing**

**Table S3. Number of reads and percentage of fragments mapped to the reference genome for CUT&Tag replicates.**

| | **reads** | **paired reads (%)** | **alignment rate (%)** |
|---|---|---|---|
| **H3K27ac replicate 1** | 20568774 | 100 | 91,25 |
| **H3K27ac replicate 2** | 14155082 | 100 | 89,06 |
| **H3K36me3 replicate 1** | 20187579 | 100 | 88,05 |
| **H3K36me3 replicate 2** | 19151354 | 100 | 80,28 |

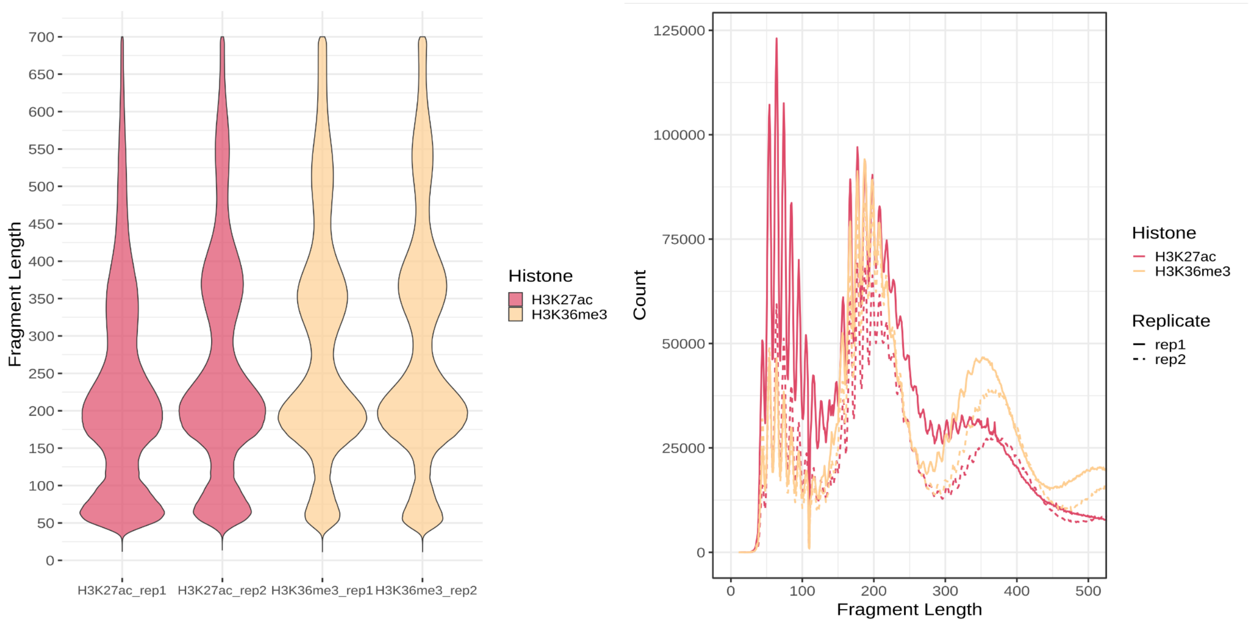


**Figure S8. Fragment length distribution and fragment length on single-base pair resolution for validating CUT&Tag results.**

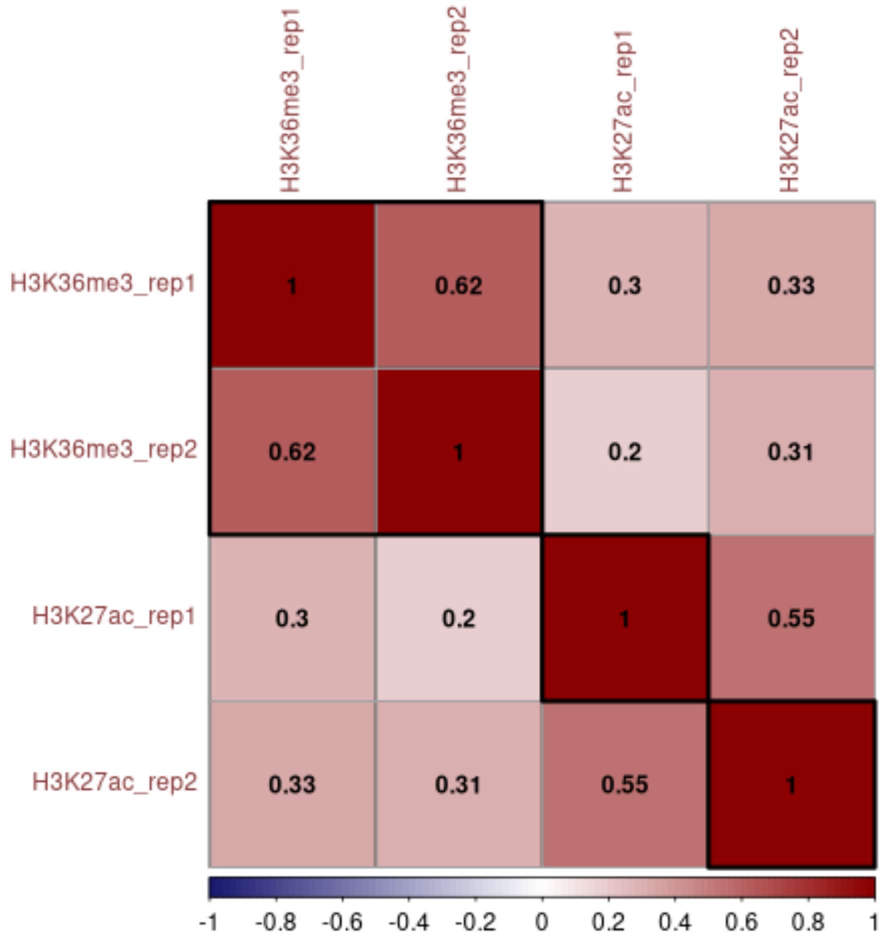


**Figure S9. Correlation matrix displaying Pearson correlation coefficients between biological replicates of embryonic histone marks.** To evaluate reproducibility, the genome was divided into 500 bp bins, and log2-transformed read counts were computed for each bin across replicates. The strength of correlation is color-coded, with darker red shades representing stronger positive correlations.

### Supplementary results

##### CpG methylation is highest in exons

**Table S4. Distribution of CpGs to different genomic features.** The last column indicates the total amount of CpGs in the respective conditions. Columns 2-6 indicate how many CpGs are covered by the respective feature. The percent value is calculated in respect to the total number.

| | Exon | Intron | 5'UTR | 3' UTR | IGR | Total |
|---|---|---|---|---|---|---|
| **Methylation ≥ 10%** | | | | | | |
| **Embryo** | 198,430 (20.7%) | 470,742 (49.1%) | 1,119 (0.1%) | 24,847 (2.6%) | 262,582 (27.4%) | 959,479 |
| **Adult** | 205,505 (20.9%) | 483,471 (49.1%) | 1,101 (0.1%) | 25,471 (2.6%) | 266,406 (27.1%) | 984,043 |
| **Methylation ≥ 90%** | | | | | | |
| **Embryo** | 26,969 (38.7%) | 29,850 (42.8%) | 161 (0.3%) | 2,593 (3.7%) | 11,807 (16.9%) | 69,747 |
| **Adult** | 7,798 (41.4%) | 7,749 (41.2%) | 50 (0.3%) | 656 (3.5%) | 3,097 (16.5%) | 18,826 |

**Table S5. Distribution of genomic features across the entire genome according to the gene annotation.**

| | Exon | Intron | IGR | Total |
|---|---|---|---|---|
| **Length in nt** | 37,881,039 (4%) | 271,949,241 (29%) | 638,956,388 (67%) | 948,786,668 |

The overlap of the 500 genes with the highest gene body methylation level per replicate was 59.8% and 58.2% for embryo and adult samples respectively. With an overlap between at least 2 samples of embryo between 8.8%-12.6% and adult 11.4%-12.4% and 15.8% - 19.6% and 17.4% - 18.4% unique genes in this category, respectively.

**Dividing annotated genes into four subsets based on methylation and expression status**

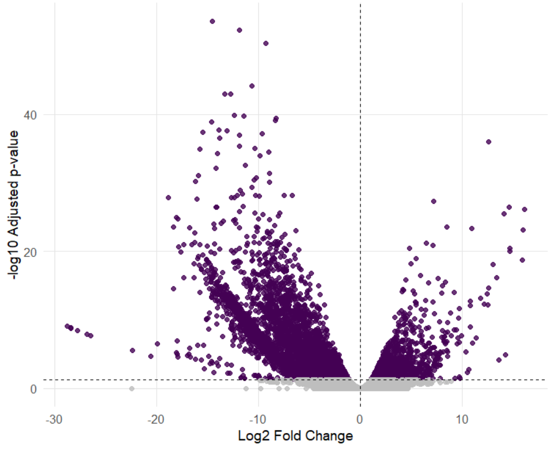


**Figure S10. Volcano plot for differentially expressed genes.** Adjusted p-value vs. the $log_2FC$ of gene expression was plotted. Dashed horizontal line indicates the significance threshold for the $log_2FC$ (1). Color indicates significant adjusted p-value. Higher expression in adults is indicated by data points on the right side.

**Table S6. Average $log_2$ fold change**

| | **average log2 fold change** | **maximum** |
|---|---|---|
| **embryo upregulated** | 6.6 | 28.8 |
| **embryo downregulated** | 3.4 | 16.2 |

For all data involving **DNA methylation** and other -omics data**,** we use a set of 25,631 genes (all genes with "NA" in any of the replicates are removed, i.e. 4,988 genes of all genes in the *L. decemlineata* annotation).
For embryo and adult samples, this set can be divided up into four subsets each.

**Table S7. Number of genes in each subset for embryo and adult.**

| | **Consolidated gene set** | **embryo** | **adult** |
|---|---|---|---|
| **Meth<10 AND FPKM≥1** | *‘not methylated / expressed‘* | 3,658 | 5,765 |
| **Meth≥10 AND FPKM≥1** | *‘methylated / expressed’* | 5,703 | 5,582 |
| **Meth<10 AND FPKM<1** | *‘not methylated / not expressed’* | 15,338 | 13,478 |
| **Meth≥10 AND FPKM<1** | *‘methylated / not expressed’* | 932 | 806 |
| **total** | | 25,631 | 25,631 |

***'methylated / expressed'*** **genes are usually longer and exclusively characterized by a drop in CpG methylation at the TSS**

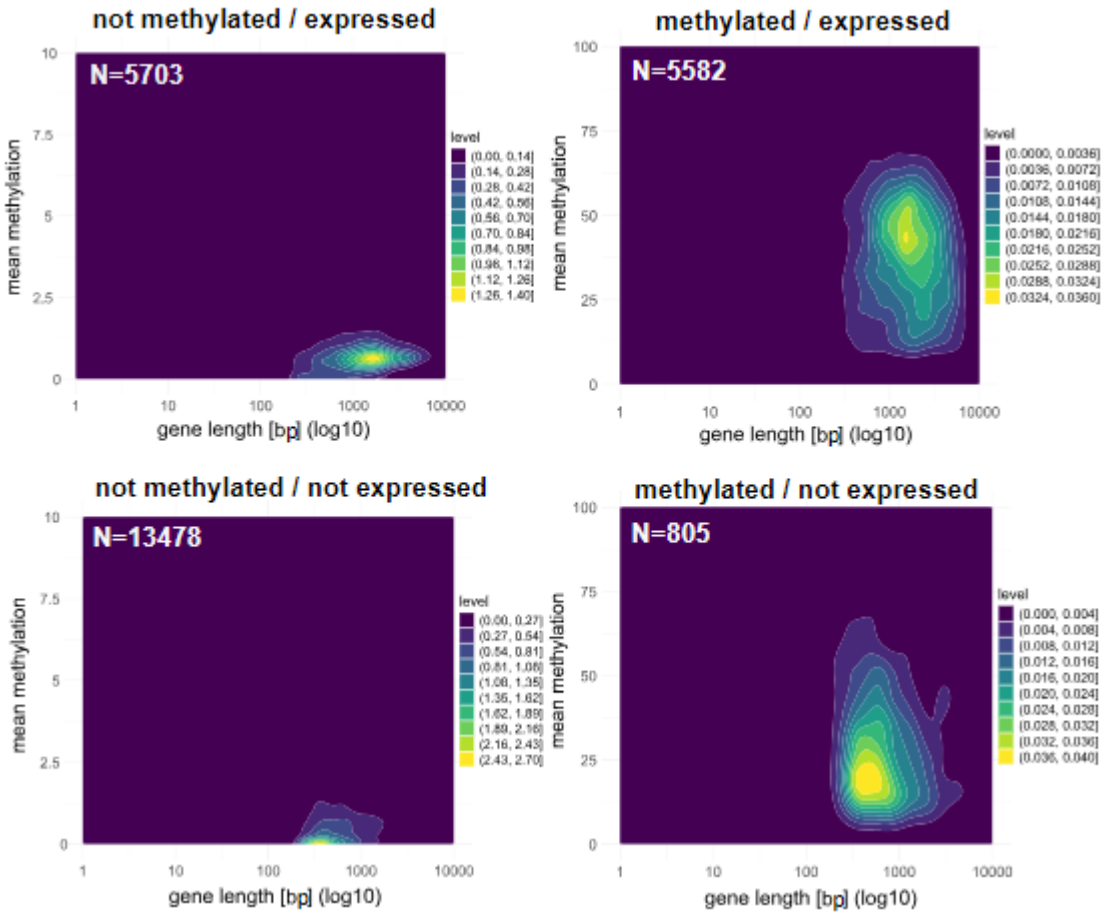


**Figure S11. Relationship between gene length and mean methylation (%) in different categories in adult *L. decemlineata*.** (Note difference in scale). The color gradient (level) represents the density of genes at different methylation levels, with brighter shades indicating regions of higher density.

**Table S8. Gene length with standard error in the four categories for embryo and adult.**

| Category | Mean gene length | Median gene length | Standard Error (SE) |
|---|---|---|---|
| **Embryo** | | | |
| ***'Not methylated / expressed'*** | 1903.68 | 1421.5 | 30.22 |
| ***'Methylated / not expressed'*** | 969.95 | 637.5 | 31.06 |
| ***'Methylated / expressed'*** | 2075.21 | 1647.0 | 21.65 |
| ***'Not methylated / not expressed'*** | 822.51 | 531.0 | 7.79 |
| **Adult** | | | |
| ***'Not methylated / expressed'*** | 1792.85 | 1369.0 | 23.18 |
| ***'Methylated / not expressed'*** | 910.60 | 603.0 | 32.96 |
| ***'Methylated /expressed'*** | 2070.68 | 1646.0 | 21.75 |
| ***'Not methylated / not expressed'*** | 718.95 | 489.0 | 6.73 |

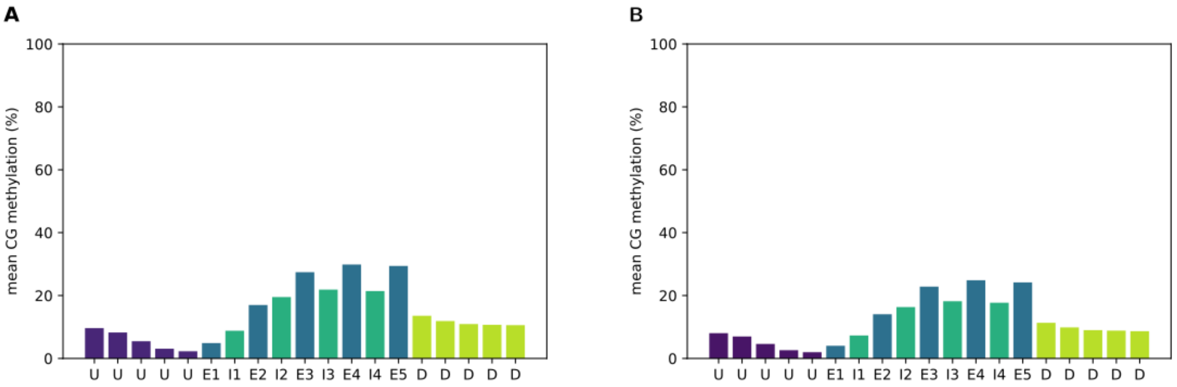


**Figure S12. Methylation level of different genic segments, in A) embryo and B) adult.** Shown is the mean methylation level of different genic segments across all annotated *L. decemlineata* genes. E1 to E5 represent the first 5 exons; I1-I4 represent the first four introns; U – upstream region; D – downstream region.

**Table S9. Amount of regions for the different categories in the embryo barplots.**

| | **All** | *'methylated / expressed'* | *'methylated / not expressed'* | *'not methylated / expressed'* | *'not methylated / not expressed'* |
|---|---|---|---|---|---|
| **UP5** | 19,875 | 4,826 | 522 | 2,955 | 9,395 |
| **UP4** | 20,048 | 4,901 | 528 | 2,954 | 9,462 |
| **UP3** | 20,134 | 4,942 | 521 | 2,947 | 9,610 |
| **UP2** | 20,329 | 4,998 | 538 | 3,009 | 9,795 |
| **UP1** | 20,689 | 5,104 | 570 | 3,192 | 10,215 |
| **E1** | 22,211 | 4,674 | 776 | 2,921 | 12,597 |
| **I1** | 13,397 | 3,413 | 259 | 3,081 | 6,517 |
| **E2** | 13,657 | 4,435 | 281 | 2,758 | 6,005 |
| **I2** | 9,863 | 3,520 | 151 | 2,568 | 3,585 |
| **E3** | 9,551 | 3,843 | 143 | 2,331 | 3,195 |
| **I3** | 7,419 | 2,957 | 94 | 2,159 | 2,200 |
| **E4** | 7,283 | 3,213 | 76 | 1,993 | 1,991 |
| **I4** | 5,897 | 2,425 | 52 | 1,786 | 1,631 |
| **E5** | 5,788 | 2,631 | 47 | 1,646 | 1,458 |
| **D1** | 12,748 | 2,625 | 388 | 1,660 | 6,801 |
| **D2** | 12,540 | 2,572 | 372 | 1,592 | 6,531 |
| **D3** | 12,426 | 2,540 | 371 | 1,571 | 6,398 |
| **D4** | 12,389 | 2,517 | 366 | 1,556 | 6,359 |
| **D5** | 12,425 | 2,513 | 367 | 1,564 | 6,314 |

**Table S10. Amount of regions for the different categories in the adult barplots.**

| | **All** | ***'methylated / expressed'*** | ***'methylated / not expressed'*** | ***'not methylated / expressed'*** | ***'not methylated / not expressed'*** |
|---|---|---|---|---|---|
| **UP5** | 20,281 | 4,859 | 530 | 2,996 | 9,592 |
| **UP4** | 20,375 | 4,922 | 519 | 2,998 | 9,650 |
| **UP3** | 20,473 | 4,941 | 529 | 3,000 | 9,795 |
| **UP2** | 20,696 | 5,001 | 544 | 3,065 | 9,996 |
| **UP1** | 21,041 | 5,088 | 572 | 3,214 | 10,418 |
| **E1** | 23,092 | 4,738 | 782 | 3,045 | 12,918 |
| **I1** | 13,697 | 3,447 | 266 | 3,108 | 6,706 |
| **E2** | 14,140 | 4,493 | 283 | 2,881 | 6,288 |
| **I2** | 9,986 | 3,555 | 156 | 2,575 | 3,653 |
| **E3** | 9,971 | 3,976 | 149 | 2,448 | 3,355 |
| **I3** | 7,556 | 3,020 | 97 | 2,176 | 2,246 |
| **E4** | 7,588 | 3,324 | 82 | 2,079 | 2,090 |
| **I4** | 6,018 | 2,496 | 53 | 1,808 | 1,656 |
| **E5** | 6,049 | 2,718 | 42 | 1,746 | 1,537 |
| **D1** | 12,974 | 2,660 | 392 | 11,671 | 6,926 |
| **D2** | 12,808 | 2,591 | 394 | 1,629 | 6,681 |
| **D3** | 12,642 | 2,560 | 383 | 1,600 | 6,518 |
| **D4** | 12,622 | 2,549 | 375 | 1,568 | 6,467 |
| **D5** | 12,693 | 2,539 | 369 | 1,574 | 6,460 |

**Gene body methylation is associated with transcription, but changes in GBM are not associated with transcription changes**

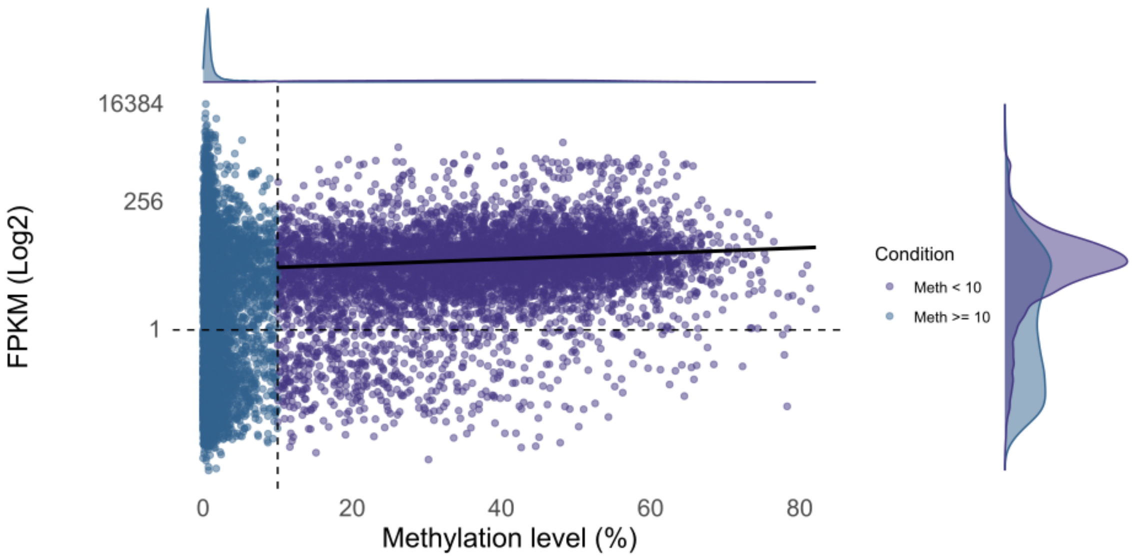


**Figure S13. Association between gene expression level (Log FPKM) and gene body methylation level in adults.** Dashed gray lines indicate thresholds used in methylation and FPKM. All values that equal 0 were removed.

For *'methylated and expressed'* genes, while the linear component of methylation is significantly positively associated with gene expression ($p < 0.001$), the quadratic term is not (embryo p-value: 0.117161; adult $p = 0.779$), indicating no curved relationship. The model explains only a small portion of the variance in gene expression ($R^2 = 0.48\%$).

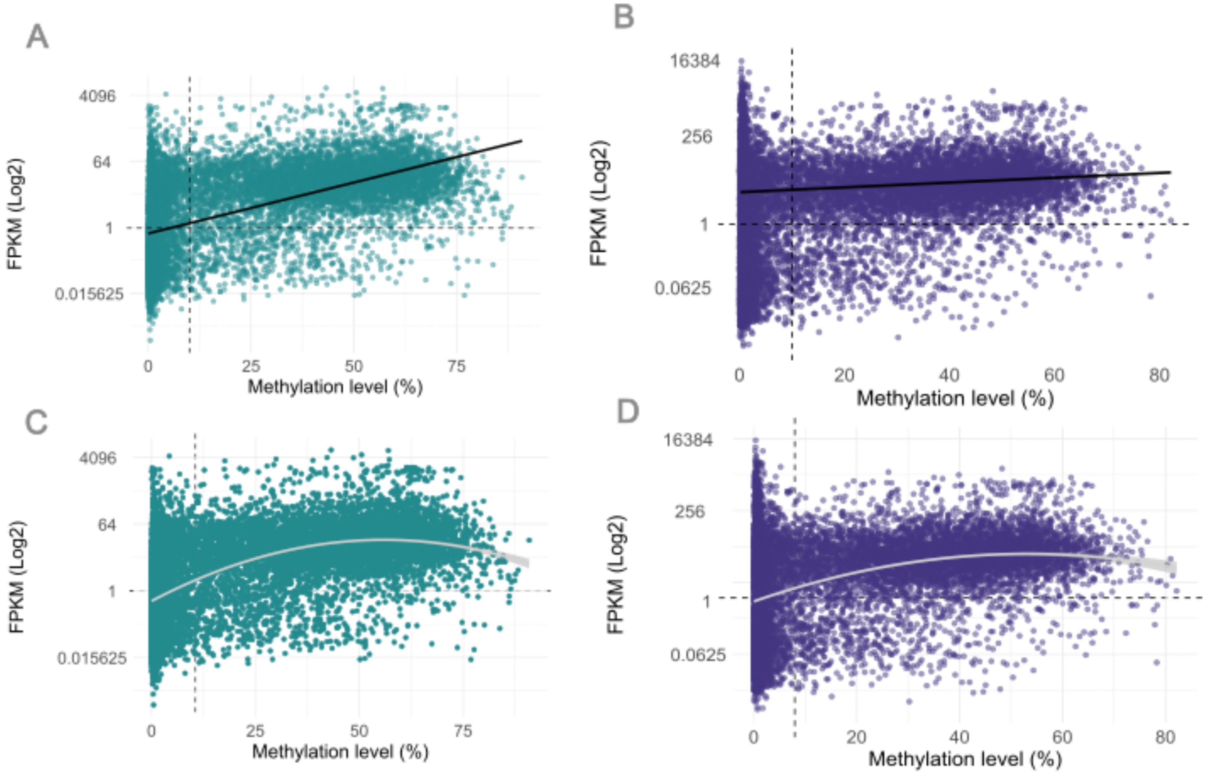


**Figure S14. Association between gene expression level (Log FPKM) and gene body methylation level in *L. decemlineata* linear and square regression.** Dashed gray lines indicate thresholds used in methylation and FPKM. Square regression was calculated for all values in A) embryo R2: 0.02534; df: 14873; p-value: 0.000591 and B) adult R2: 0.00066; df: 16197; p-value 0.001753 in C) embryo R2: 0.002757; df: 5700; p-value: 0.117161 n.s. and D) adult R2: 0.00444; df: 5579; p-value: 0.779 n.s.

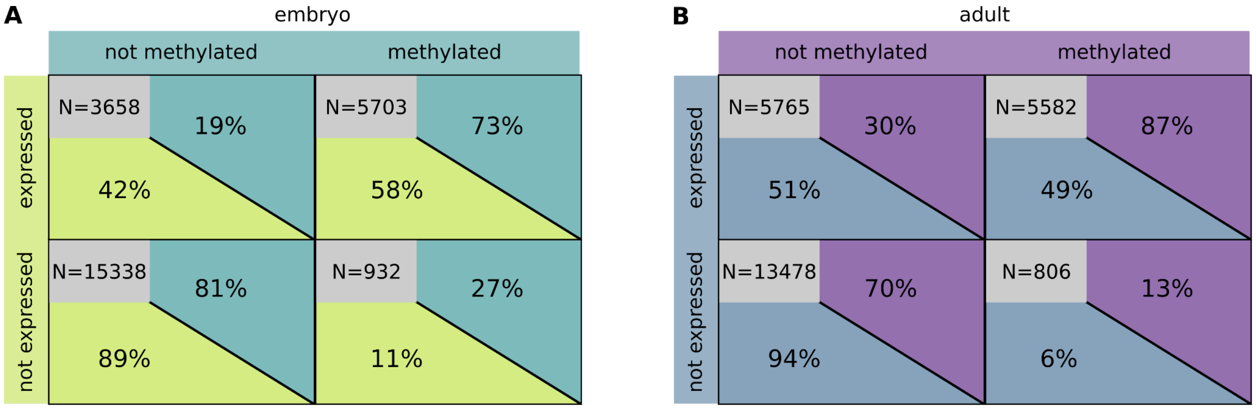


**Figure S15. Matrix summarizing the relationship between methylation and expression status in embryos and adults.** (Example upper left corner (embryo): 42% of expressed genes are not methylated, while only 19% of not methylated genes are expressed.)

**Table S11. Genes changing category assignment from embryo to adult given in absolute numbers.**

| | *'not methylated / expressed'* | *'methylated / expressed'* | *'not methylated / not expressed'* | *'methylated / not expressed'* | |
|---|---|---|---|---|---|
| *'not methylated / expressed'* | 2906 | 6 | 745 | 1 | 3658 |
| *'methylated / expressed'* | 127 | 5454 | 12 | 110 | 5703 |
| *'not methylated / not expressed'* | 2705 | 1 | 12610 | 22 | 15338 |
| *'methylated / not expressed'* | 27 | 121 | 111 | 673 | 932 |
| | 5765 | 5582 | 13478 | 806 | |

**Table S12. Percentage of embryonic genes changing category assignment from embryo to adult.**

| embryo/adult | *'not methylated / expressed'* | *'methylated / expressed'* | *'not methylated / not expressed'* | *'methylated / not expressed'* | |
|---|---|---|---|---|---|
| *'not methylated / expressed'* | 79.4% | 0.2% | 20.4% | 0.0% | 100% |
| *'methylated / expressed'* | 2.2% | 95.7% | 0.2% | 1.9% | 100% |
| *'not methylated / not expressed'* | 17.6% | 0.0% | 79.3% | 0.1% | 100% |
| *'methylated / not expressed'* | 2.9% | 13.0% | 11.9% | 72.2% | 100% |

**Gene Ontology enrichment indicates functional differences of methylated and not methylated genes**

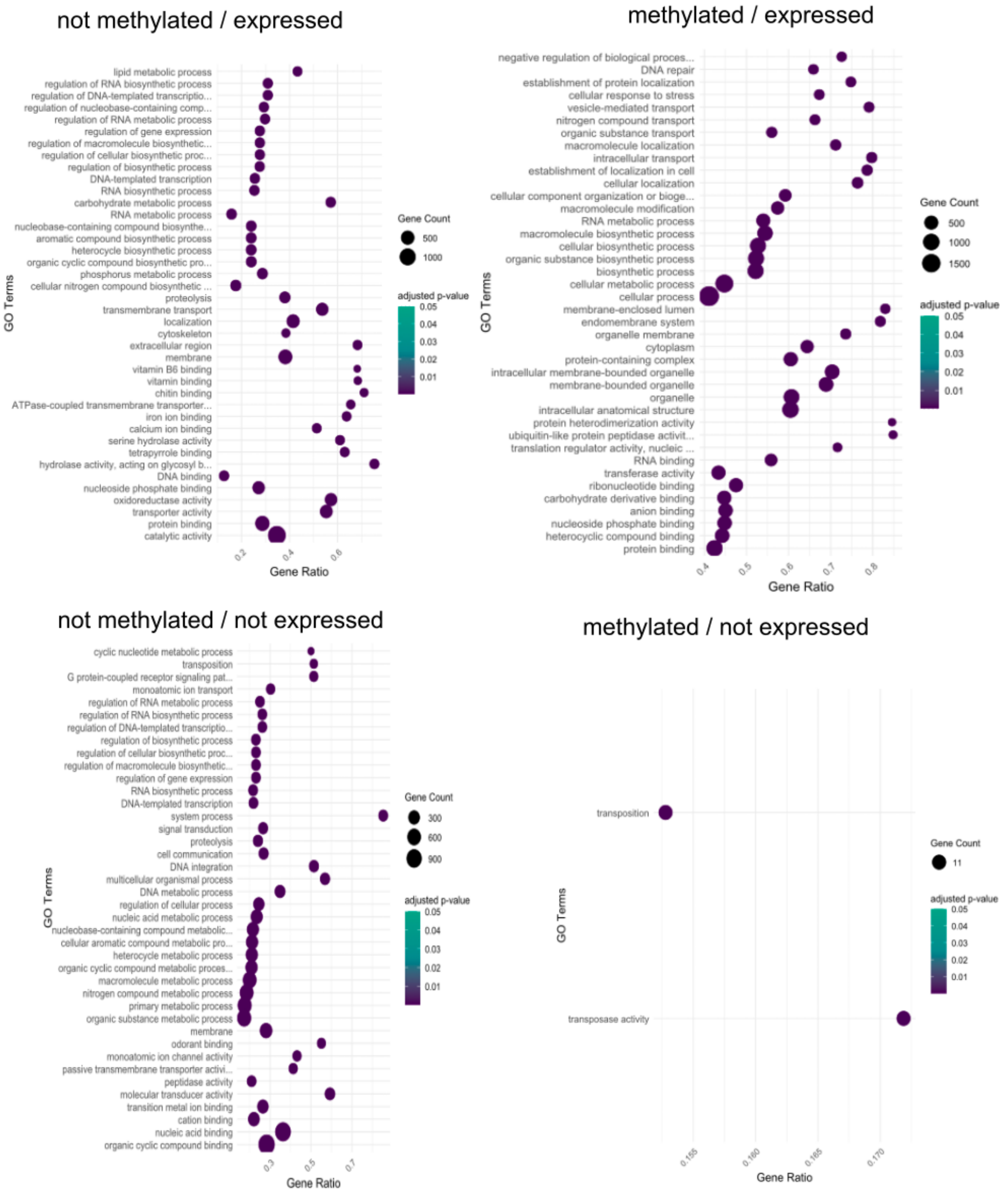


**Figure S16. Adult top 40 significantly enriched Gene Ontology (GO) terms for the four subsets.** The p-value was adjusted using the Benjamini-Hochberg procedure. Gene count is the number of genes associated with a GO term, whereas 'Gene Ratio' is the percentage of genes of the specific subset in the given GO terms. Categories vary in their number of genes included, resulting in different numbers of enriched GO terms. (*'not methylated / expressed'* = 135 GO; *'methylated / expressed*' = 123 GO; *'not methylated / not expressed*' = 120 GO; '*methylated / not expressed*' = 2 GO).

**Only H3K36me3 is associated with CpG methylation, while H3K27ac and H3K36me3 are associated with active transcription**

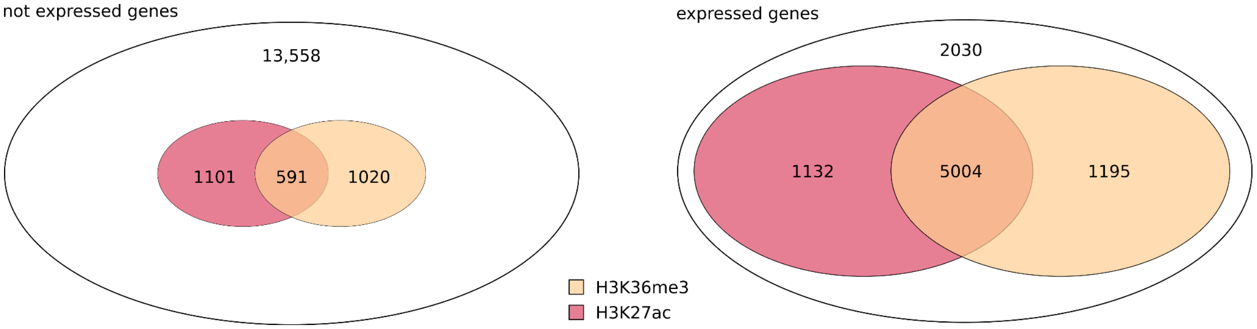


**Figure S17. A) Venn diagram showing the proportion of expressed genes with enrichment of H3K27ac, H3K36me3 or both.** Total number of expressed genes = 9,361; percentage of genes carrying at least one histone mark: 78.31%; percentage of genes with H3K27ac enrichment: 65.53%; percentage of genes with H3K36me3 enrichment: 66.22%; percentage of genes with enrichment of both marks: 53.45%. **B) Venn diagram showing the proportion of not expressed genes with enrichment of H3K27ac, H3K36me3 or both.** Total number of expressed genes = 16,270; percentage of genes carrying at least one histone mark: 16.66%; percentage of genes with H3K27ac enrichment: 10.39%; percentage of genes with H3K36me3 enrichment: 9.9%; percentage of genes with enrichment of both marks: 3.63%.

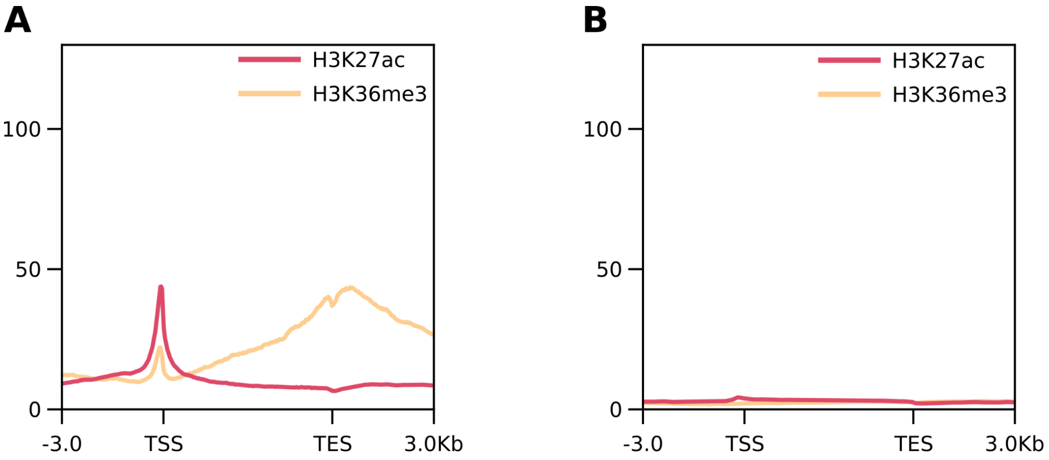


**Figure S18. Enrichment patterns of H3K27ac and H3K36me3 in *'expressed'* (A) and *'not expressed'* (B) genes.** Shown are the profiles for replicate 2.

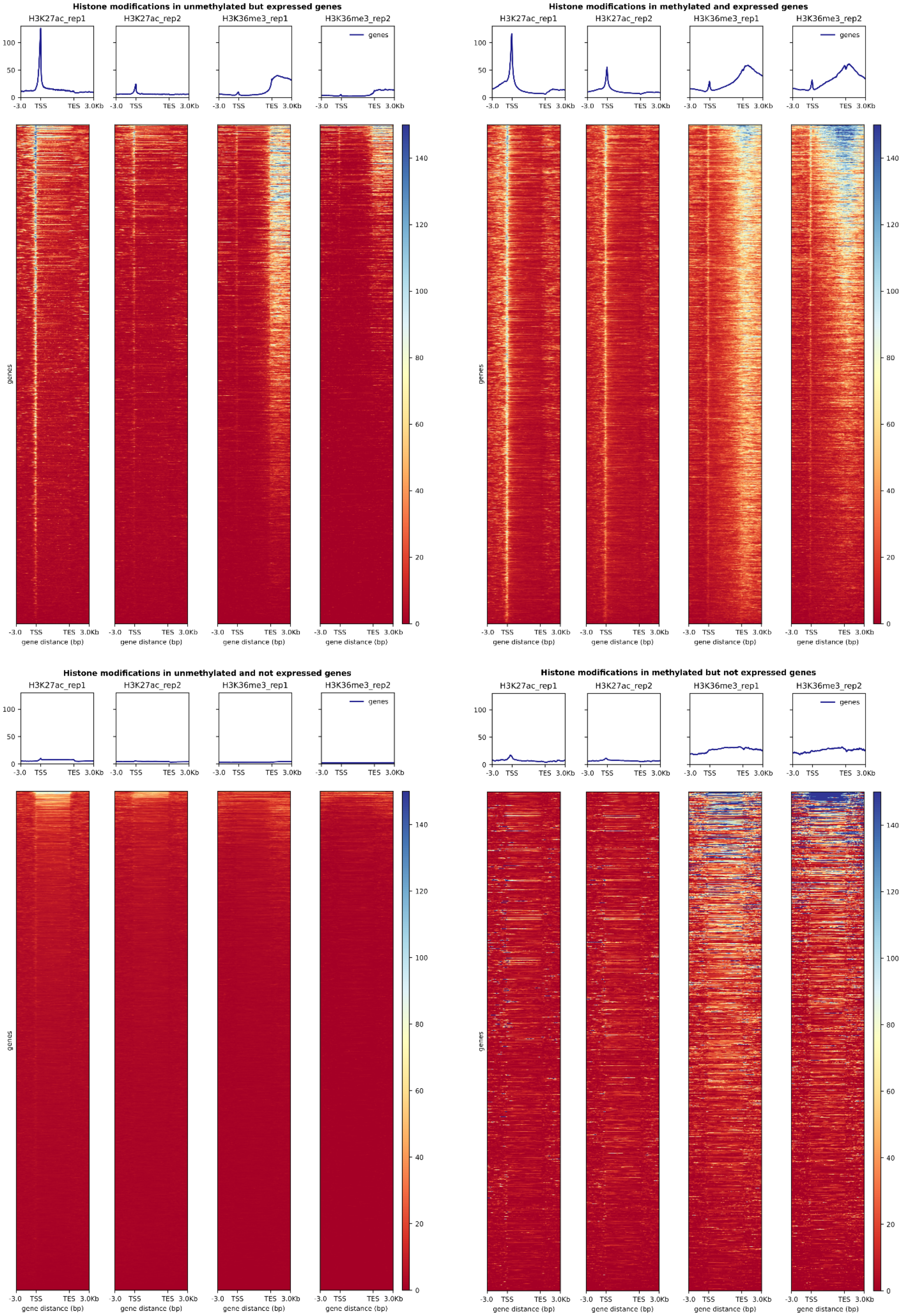


**Figure S19. Profiles and corresponding heatmaps of H3K27ac and H3K36me3 enrichment in gene bodies for each replicate.** Genes are normalized to a length of 5 kb, with 3 kb upstream and downstream of TSS and TES, respectively.
